# The Interferon-inducible NAMPT acts as a protein phosphoribosylase to restrict viral infection

**DOI:** 10.1101/2023.10.12.562112

**Authors:** Shu Zhang, Na Xie, Yongzhen Liu, Chao Qin, Ali Can Savas, Ting-Yu Wang, Shutong Li, Youliang Rao, Alexandra Shambayate, Tsui-Fen Chou, Charles Brenner, Canhua Huang, Pinghui Feng

## Abstract

As obligate intracellular pathogens, viruses often activate host metabolic enzymes to supply intermediates that support progeny production. Nicotinamide phosphoribosyltransferase (NAMPT), the rate-limiting enzyme of the salvage NAD^+^ synthesis, is an interferon-inducible protein that inhibits the replication of several RNA and DNA viruses with unknown mechanism. Here we report that NAMPT restricts herpes simplex virus 1 (HSV-1) replication via phosphoribosyl-hydrolase activity toward key viral structural proteins, independent of NAD^+^ synthesis. Deep mining of enriched phosphopeptides of HSV-1-infected cells identified phosphoribosylated viral structural proteins, particularly glycoproteins and tegument proteins. Indeed, NAMPT de-phosphoribosylates viral proteins *in vitro* and in cells. Chimeric and recombinant HSV-1 carrying phosphoribosylation-resistant mutations show that phosphoribosylation promotes the incorporation of structural proteins into HSV-1 virions and subsequent virus entry. Moreover, loss of NAMPT renders mice highly susceptible to HSV-1 infection. The work describes a hidden enzyme activity of a metabolic enzyme in viral infection and host defense, offering a system to interrogate roles of phosphoribosylation in metazoans.

## INTRODUCTION

Viruses are obligate intracellular pathogens and rely on cellular machinery for the synthesis of diverse macromolecules that collectively support viral replication. Not surprisingly, viruses often activate cellular metabolic enzymes to fuel viral replication presumably via providing metabolites for the synthesis of viral structural building blocks.^1^ Indeed, interferon-inducible enzymes that interfere with metabolites essential for viral replication can function to restrict the infection of particular viruses, thus acting as effectors within the broad IFN response network.^2,3^ For example, SAMHD1 hydrolyzes deoxyribonucleosides triphosphate (dNTP) to limit HIV replication in nonpermissive cells, while viperin catalyzes the synthesis of 3’-deoxy-3’,4’-didehydro-cytidine triphosphate that terminates the RNA-dependent RNA polymerization of diverse RNA viruses.^4,5^ Cholestrol-25-hydroxylase (CH25H) converts cholesterol into 25-hydroxylcholesterol that inhibits the membrane fusion between cells and diverse envelope viruses.^6^ These studies illustrate a paradigm that interferon-inducible metabolic enzymes restrict viral replication via their metabolites. Whether and how IFN-inducible metabolic enzymes limit viral infection independent of their metabolites are poorly understood.

Nicotinamide adenine dinucleotide (NAD^+^) is a vital enzyme cofactor in all living cells.^7^ NAD^+^ homeostasis is maintained by a balance between its synthesis and consumption. NAD^+^ is consumed chiefly by NAD^+^-dependent enzymes, such as polyADP-ribose polymerases (PARPs) and sirtuin deacetylases (SIRTs), and by glycohydrolases such as CD38 and sterile alpha and toll/interleukin receptor motif-containing protein 1 (SARM1), yielding nicotinamide (NAM) as a byproduct.^8,9^ Protein ADP/polyADP-ribosylation plays critical roles in diverse biological processes such as gene expression, DNA damage response, and translation.^10–12^ Although phosphoribose of proteins can be processed from ADP/polyADP-ribose by nudix hydrolase 16 (NUDT16), ectonucleotide pyrophosphatase/phosphodiesterase 1 (ENPP1), or snake venom phosphodiesterase (svPDE) in vitro,^13,14^ the biogenesis and roles of phosphoribosyl moieties on proteins are poorly understood in metazoans.^15^ NAM is converted into nicotinamide mononucleotide (NMN) by NAM phosphoribosyltransferase (NAMPT), the rate-limiting enzyme of the salvage NAD^+^ synthesis, to regenerate NAD^+^. Pivotal roles of NAMPT in diverse biological processes are reflected by its upregulation in microbial infection and cancer cells, as well as its deficiency in neurodegenerative diseases.^9,16,17^ Previous studies have implicated NAMPT in antiviral defense against diverse viruses, including HIV, West Nile virus, Venezuelan equine encephalitis virus.^18^ It was suggested that NAMPT provides NAD^+^ in immune cells to fend off the infection of these viral pathogens. Here we describe that NAMPT exerts antiviral activity against herpes simplex virus 1 (HSV-1) independent of its metabolites. Mechanistically, NAMPT directly de-phosphoribosylates viral structural proteins, thereby impeding HSV-1 virion assembly and entry of subsequent infection.

## RESULTS

### NAMPT restricts HSV-1 lytic replication independent of NAD^+^

By analyzing metabolites of HSV-1-infected cells, we found that HSV-1 significantly depletes NAD^+^ and its related metabolites, such as NADH, NADP, and NAM, during lytic replication (Figure 1A and S1A). In mammalian cells, NAD^+^ is often rapidly replenished via the activity of the rate-limiting NAMPT of the salvage pathway (Figure 1B). Thus, we depleted key enzymes responsible for NAD^+^ synthesis, including NAMPT, nicotinamide riboside kinase (NRK), NMN adenylyltransferase 1 (NMNAT1), and NAD^+^ synthetase (NADSYN), via shRNA-mediated knockdown and examined HSV-1 lytic replication (Figure 1B). Surprisingly, depletion of NAMPT, but not the other three enzymes, increased the extracellular HSV-1 titer (Figure 1C and 1D). Transient depletion of all four enzymes had no effect on cell viability (Figure S1B). Similarly, depletion of NAMPT also increased HSV-1 lytic replication in HeLa cells, as shown by extracellular and intracellular HSV-1 titers (Figure 1E). These results suggest that NAMPT is antiviral against HSV-1.

**Figure 1.**
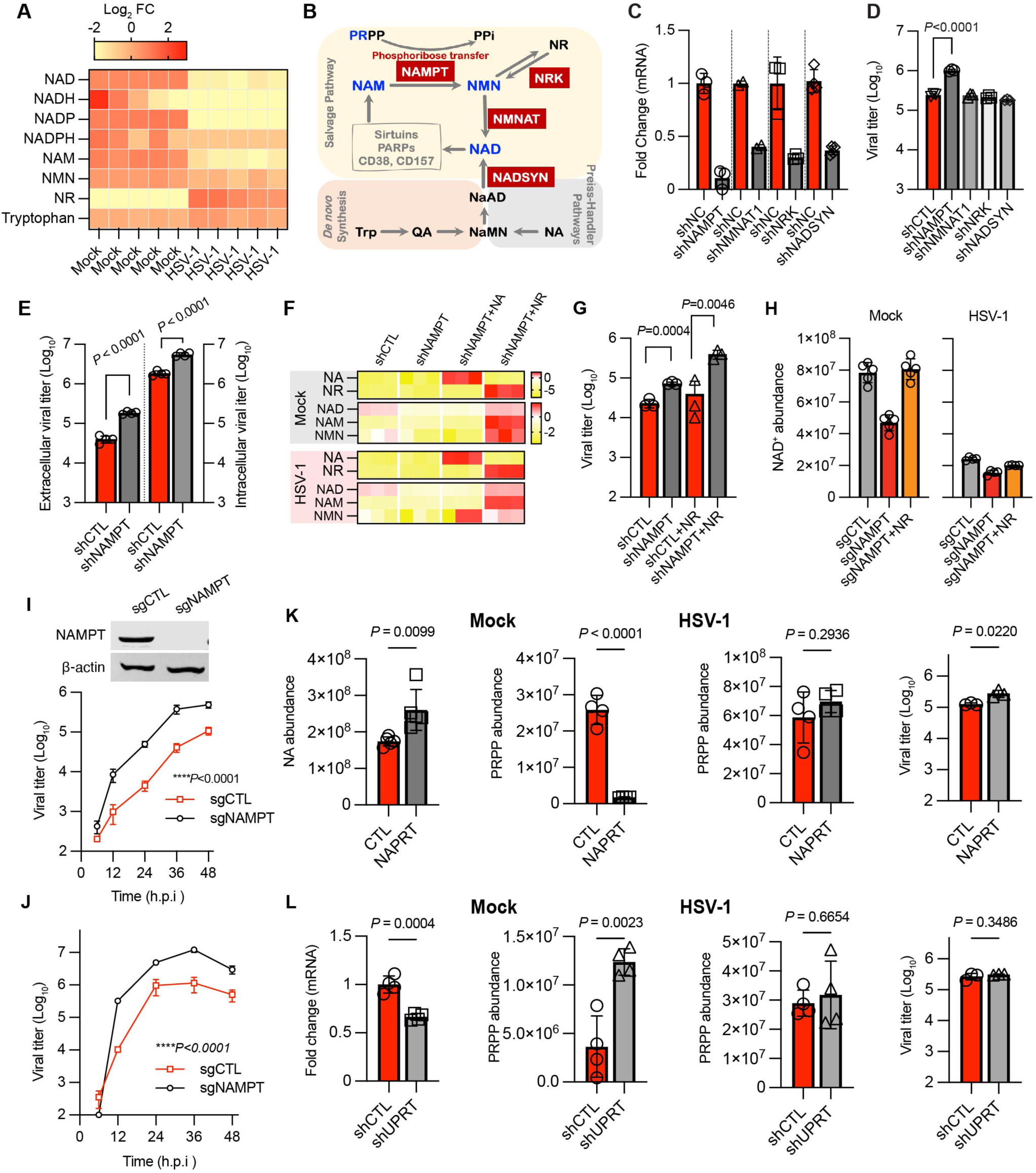
NAMPT restricts HSV-1 lytic replication. (A) Heatmap of the NAD^+^-related metabolites in HSV-1-infected HepG2 cells. (B) Schematic illustration of the NAD^+^ synthesis pathways. (C) Knockdown of NAMPT, NMNAT1, NRK and NADSYN in 293T cells validated via quantitative RT-PCR. (D) HSV-1 titer in the medium of 293T cells with depletion in NAMPT, NMNAT1, NRK and NADSYN. (E) Extracellular and intracellular HSV-1 titers in control (shCTL) and NAMPT-depleted (shNAMPT) HeLa cells at 48 h post-infection (MOI=0.1). (F) Heatmap of the NAD^+^-related metabolites in mock- and HSV-1 infected HeLa cells, supplemented with nicotinic acid (NA, 0.1 mM) or nicotinamide riboside (NR, 0.1 mM). (G) HSV-1 titer in control and shNAMPT HeLa cells without or with supplementation of NR (0.1 mM) at 48 h.p.i. (MOI=0.1). (H) NR (0.1 mM) restores NAD^+^ in mock- and HSV-1-infected shNAMPT HeLa cells. (I and J) Growth curve of HSV-1 is determined by plaque assay using medium (I) and cells with three rounds of freeze/thaw (J) in control (sgCTL) and NAMPT-knockout (sgNAMPT) HeLa cells. (K and L) HeLa cells were infected with lentivirus expressing NAPRT (K) or shUPRT (L). HeLa cells were mock- or HSV-1-infected as indicated in (A). Intracellular PRPP was determined by LC-MS at 12 h.p.i. and viral titer in the medium was determined by plaque assay using Vero monolayer. Statistical significance was calculated using unpaired two-tailed Student’s *t*-tests for D, E, G, K and L; two-way ANOVA for I and J.

NAMPT is crucial for intracellular NAD^+^ regeneration. Indeed, mass spectrometry analysis showed that NAMPT depletion reduced intracellular NAD^+^ and NMN in mock- and HSV-1-infected HeLa cells by >50% (Figure S1C). Addition of NR, but not nicotinic acid (NA), increased intracellular NMN and NAD^+^ levels in a dose-dependent manner (Figure 1F, S1C, and S1D). The dose-dependent effect of exogenous NR to restore intracellular NAD^+^ was confirmed in mock- and HSV-1-infected sgNAMPT HeLa cells (Figure S1E and S1F). The increase in intracellular NR and NMN by NR supplementation was more pronounced in HSV-1-infected cells than in mock-infected cells. NR restored intracellular NAD^+^ of NAMPT-knockout cells to that of wildtype HeLa cells at 0.1 mM (Figure S1F). We then supplemented NAMPT-depleted HeLa cells with NR (0.1 mM) and found that addition of NR further boosted HSV-1 lytic replication in NAMPT-depleted HeLa cells (Figure 1G and 1H). Similarly, NAMPT knockdown increased HSV-1 lytic replication in human HepG2 hepatocarcinoma cells (Figure S1G). Furthermore, the expression of exogenous NAMPT in 293T cells reduced HSV-1 replication, as indicated by extracellular infectious virions, in a dose-dependent manner (Figure S1H). Finally, the depletion of NAMPT in mouse embryonic fibroblasts (MEFs) and human foreskin fibroblasts (HFF), primary cells from mice and humans, significantly increased HSV-1 lytic replication, as determined by the extracellular HSV-1 (Figure S1I and S1J). Thus, NAMPT reduces HSV-1 lytic replication independent of its activity in NAD^+^ synthesis.

### Antiviral activity of NAMPT is independent of PRPP in HSV-1 infection

In addition to salvaging NAD^+^ synthesis, NAMPT consumes phosphoribosyl-pyrophosphate (PRPP), which may be a limiting factor for HSV-1 replication. We then manipulated intracellular PRPP levels by over-expressing nicotinic acid phosphoribosyltransferase (NAPRT) or silencing uracil phosphoribosyltransferase (UPRT, also known as UMP synthetase) and examined HSV-1 lytic replication. NAPRT catalyzes the Preiss-Handler pathway using nicotinic acid and PRPP as substrates, while UPRT adds phosphoribosyl moiety to uracil to synthesize UMP. Thus, NAPRT overexpression and UPRT depletion are expected to decrease or increase intracellular PRPP levels, respectively. Indeed, NAPRT expression decreased PRPP by ∼90% in mock-infected HeLa cells, while it had no effect on HSV-1 replication (Figure 1H). On the other hand, UPRT depletion at mRNA level by ∼35% increased PRPP three-fold and demonstrated no effect on HSV-1 replication (Figure 1I). Thus, NAMPT restricts HSV-1 lytic replication independent of intracellular NAD^+^ and PRPP levels.

### NAMPT is antiviral against HSV-1 in mice

To probe the *in vivo* role of NAMPT in HSV-1 infection, we induced NAMPT deletion with tamoxifen in conditional NAMPT-knockout (KO) mice (*Nampt^flox/flox^:Cre*) that were bred from *Nampt^tm10leo^*/ImaiJ and B6;129-*Gt(ROSA)26Sor^tm1(cre/ERT2)Tyj^*/J. When the mice were treated with tamoxifen, we found that NAMPT depletion was apparent in the liver and spleen, while not detectable in the brain, kidney, heart, and lung (Figure 2A and S2A). At the time of HSV-1 infection, wildtype and NAMPT-KO mice demonstrated no significant difference in body weight (Figure S2B). When mice were infected with a sub-lethal dose of HSV-1 (5×10^7 PFU) via intravenous injection that enables synchronized and systemic infection, we found that 90% of NAMPT-KO mice succumbed to HSV-1 infection by day 6 post-infection, while 70% of wildtype mice survived HSV-1 infection under the same conditions (Figure 2B). Consistent with this observation, hematoxylin & eosin (H&E) staining revealed localized tissue destruction throughout the liver of NAMPT-KO mice, but not in that of wildtype mice (Figure 2C and S2C). Immunohistochemical staining showed that HSV-1 lytic antigens, including UL37 and UL47, were readily detected in areas with tissue destruction (Figure 2C and S2C). The HSV-1 load in the livers of NAMPT-KO mice was four orders of magnitude higher in viral titer and 2 orders of magnitude hinger in genome copy number than that in wildtype mice (Figure 2D and 2E). Taken together, NAMPT demonstrates antiviral activity against HSV-1 in mice.

**Figure 2.**
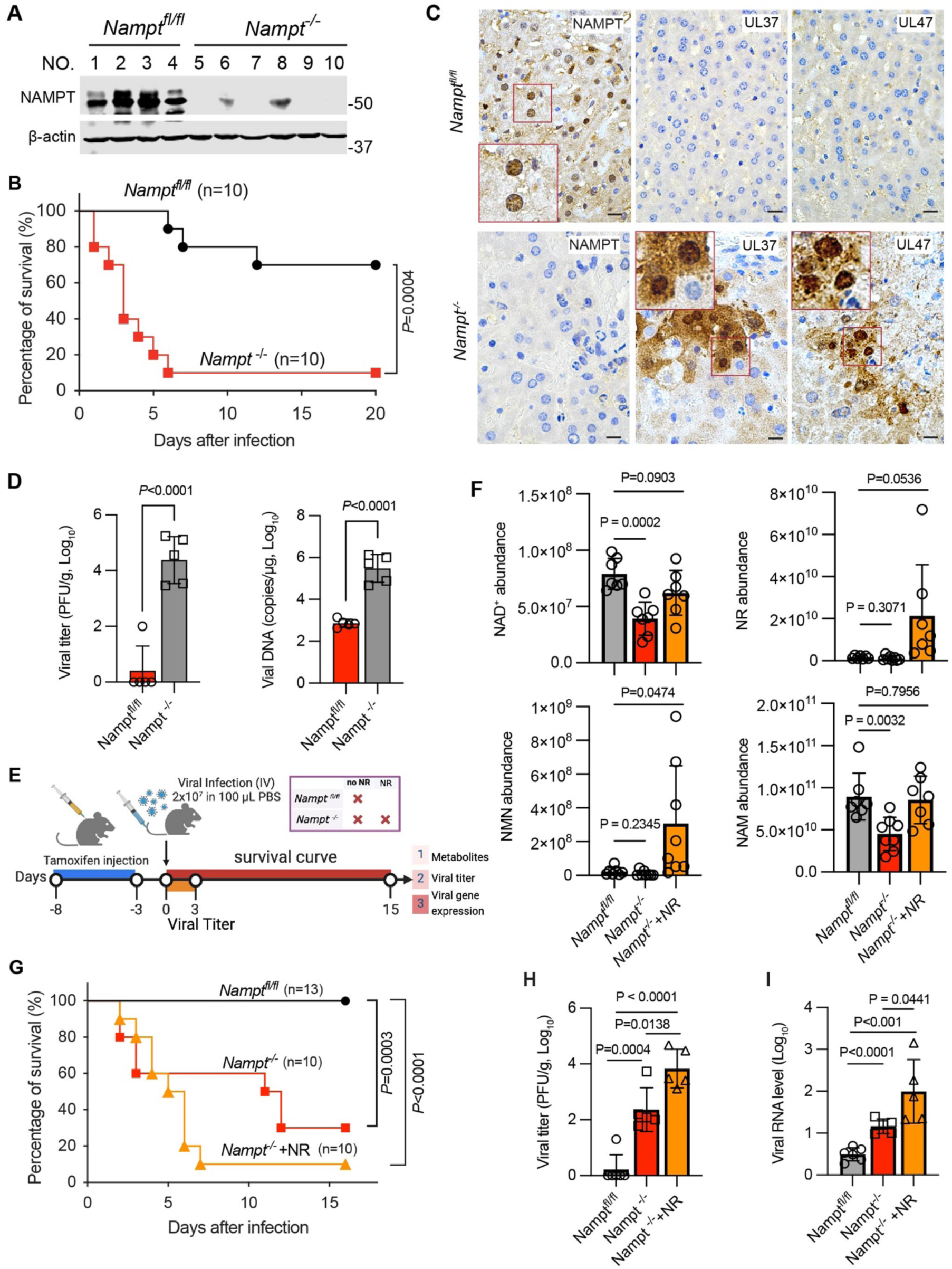
NAMPT restricts HSV-1 lytic replication in mice. (A) Immunoblotting analysis of the liver of *Nampt^fl/fl^* and *Nampt^-/-^*mice infected with HSV-1, with indicated antibodies. (B to D) Age (12-14-week-old)- and gender-matched *Nampt^fl/fl^* and *Nampt^-/-^* mice were infected with HSV-1 (5x10^7^ PFU, intraperitoneal injection) and mouse survival was recorded (B), immunohistology was performed with antibodies against NAMPT and HSV-1 antigens (UL37 and UL47)(C), HSV-1 load was analyzed by plaque assays using liver homogenates and viral genome copy number by real-time PCR using extracted total DNA (D). (E to I) Diagram of the experimental design involving NAMPT deletion via tamoxifen injection, HSV-1 infection, mouse survival, HSV-1 replication, and host immune response (E). NR was injected (400 mg/kg/day) intraperitoneally at the time of tamoxifen induction. Levels of the NAD^+^-related metabolites (F) in the liver of wildtype, NAMPT-knockout and NAMPT-knockout with NR supplementation were determined immediately before HSV-1 infection. Mouse survival was recorded after HSV-1 infection (2x10^7^ PFU, intravenous) (G), and HSV-1 replication was determined by plaque assay (H) and viral gene expression (I). Scale bar, 10 μm. Statistical significance was calculated using the log-rank test for mouse survival data (B and G), unpaired two-tailed Student’s *t*-tests for D, F, H and I.

To assess whether exogenous NR increases HSV-1 replication in mice as it did in cultured cells, we performed NR intraperitoneal injections (400 mg/kg per day) in NAMPT-knockout mice starting at the time of tamoxifen induction. We then examined mouse survival, NAD^+^-related metabolites, and HSV-1 replication in the liver where NAMPT had been depleted (Figure 2F). LC-MS analysis showed that loss of NAMPT reduced NAD^+^ levels by ∼50% in the liver. This reduction was restored to the level found in the livers of wildtype mice by exogenous NR administration, and these mice had similar body weights (Figure S2D and S2E). Considering that NR may increase HSV-1 replication, we infected mice with a dose (2×10^7 PFU) lower than the sublethal dose of HSV-1 and monitored mouse survival. We found that 40% and 70% of NAMPT-KO mice succumbed to HSV-1 infection by day 4 and 12, respectively, while no wildtype mice died from HSV-1 infection (Figure 2G). Remarkably, 90% of NAMPT-KO mice that received NR injection succumbed to HSV-1 infection by day 7. The plaque assay showed that the loss of NAMPT increased HSV-1 lytic replication by two orders of magnitude, and this effect was further elevated by ∼50-fold with NR supplementation (Figure 2H). The increased HSV-1 titer also correlated with a similar pattern of viral lytic gene expression, as analyzed by real-time PCR (Figure S2F). These results collectively show that NR supplementation further increases HSV-1 lytic replication in NAMPT-KO mice, thus NAMPT restricts HSV-1 infection independent of NAD^+^ synthesis.

### Loss of NAMPT does not compromise innate immune defense against HSV-1

Immune defense is critical for HSV-1 lytic replication and NAMPT depletion may compromise host antiviral immune response against HSV-1. Considering the rapid infection of HSV-1 in mice, we examined innate immune response in wildtype and NAMPT-KO cells and mice. To determine whether the loss of NAMPT compromises host antiviral immune defense, we examined the expression of inflammatory genes in NAMPT-deleted cultured cells and mice upon HSV-1 infection. We found that NAMPT deletion in HeLa and HepG2 cells did not significantly decrease the expression of inflammatory genes, represented by *IFNb*, *ISGs*, and *IRF7* (Figure S3A-S3C). NR supplementation to sgNAMPT HepG2 cells had no significant effect on the expression of inflammatory genes (Figure S3C). Furthermore, real-time PCR analysis showed that, in response to HSV-1 infection, most of inflammatory genes, such as *Ifnb*, *Tnf*, *Isg*s, and *Il6*, were expressed without significant difference in the livers of wildtype and NAMPT-KO mice, both without or with NR supplementation. Although *Ifnb* and *Isg56* were expressed at slightly lower levels in mock-infected NAMPT-KO mice regardless of NR supplementation, the induction was minimal, and the difference was diminished by HSV-1 infection (Figure S3D). Interestingly, the expression of two chemokines, *Cxcl1* and *Cxcl2* (*Mip2*), was significantly higher in the livers of NAMPT-KO and NAMPT-KO supplemented with NR than in wildtype mice with HSV-1 infection. This pattern was similar to the increased expression of *Cxcl1* observed even without HSV-1 infection (Figure S3D). These results collectively support the conclusion that loss of NAMPT does not compromise host antiviral innate immune response against HSV-1.

### Evidence that NAMPT acts on HSV-1 tegumentation in the TGN

NAMPT localizes to diverse subcellular compartments within mammalian cells, including the cytosol, nucleus, mitochondrion, and endoplasmic reticulum (ER)/trans-Golgi network (TGN).^19^ To enable NAMPT detection by immunofluorescence and immunogold staining, we utilized sgNAMPT HeLa cells that were reconstituted with HA-NAMPT expression. Upon HSV-1 infection, we observed redistribution of NAMPT with the progression of HSV-1 lytic replication. Specifically, NAMPT was enriched and localized to the peri-nuclear region as punctate structures at late stages of infection. Further immunofluorescence staining showed that these punctate structures were positive for NAMPT, UL19, and TGN46 (Figure 3A). These vesicular structures extended to regions proximal to the plasma membrane, suggesting that these vesicles containing HSV-1 virions were likely en route to the plasma membrane for release. NAMPT was also observed to colocalize with gB and UL19 in structures reminiscent of the TGN, which is the compartment that glycoproteins and tegument proteins are packaged into virion particles (Figure S4A).^20^ We further examined the interaction of NAMPT with HSV-1 assembly and egress processes by electron microscopy. In subcellular compartments that were positive for TGN46, double-membrane structures containing HSV-1 virions were observed and NAMPT localized to the outer membrane (Figure 3B, left top panel). An invaginating virion within the TGN was detected through immunostaining, revealing that NAMPT and TGN46 were found at two separate ends of the unclosed membrane structure (Figure 3B, left bottom panel). NAMPT was also detected in proximity to virions within the lumen of the double-layered membrane structure, which is localized in the vicinity of TGN stacks (Figure 3B, right panel, S4B, and S4C). As shown in Figure 3C, the presence of HSV-1 virions in the lumen of the TGN was further confirmed through staining for UL19 and NAMPT. Within a subcellular compartment where a vesicle containing HSV-1 virions was pinching from the membrane, abundant NAMPT was detected at the site of membrane closure (Figure 3C, left panel). This observation provides the temporal and spatial information regarding the action of NAMPT during HSV-1 lytic replication. We also detected subcellular regions that were stained predominantly for NAMPT and UL19 (Figure 3C, right panel). These results collectively support the conclusion that NAMPT acts within the proximity of the TGN to restrict HSV-1 progeny assembly.

**Figure 3.**
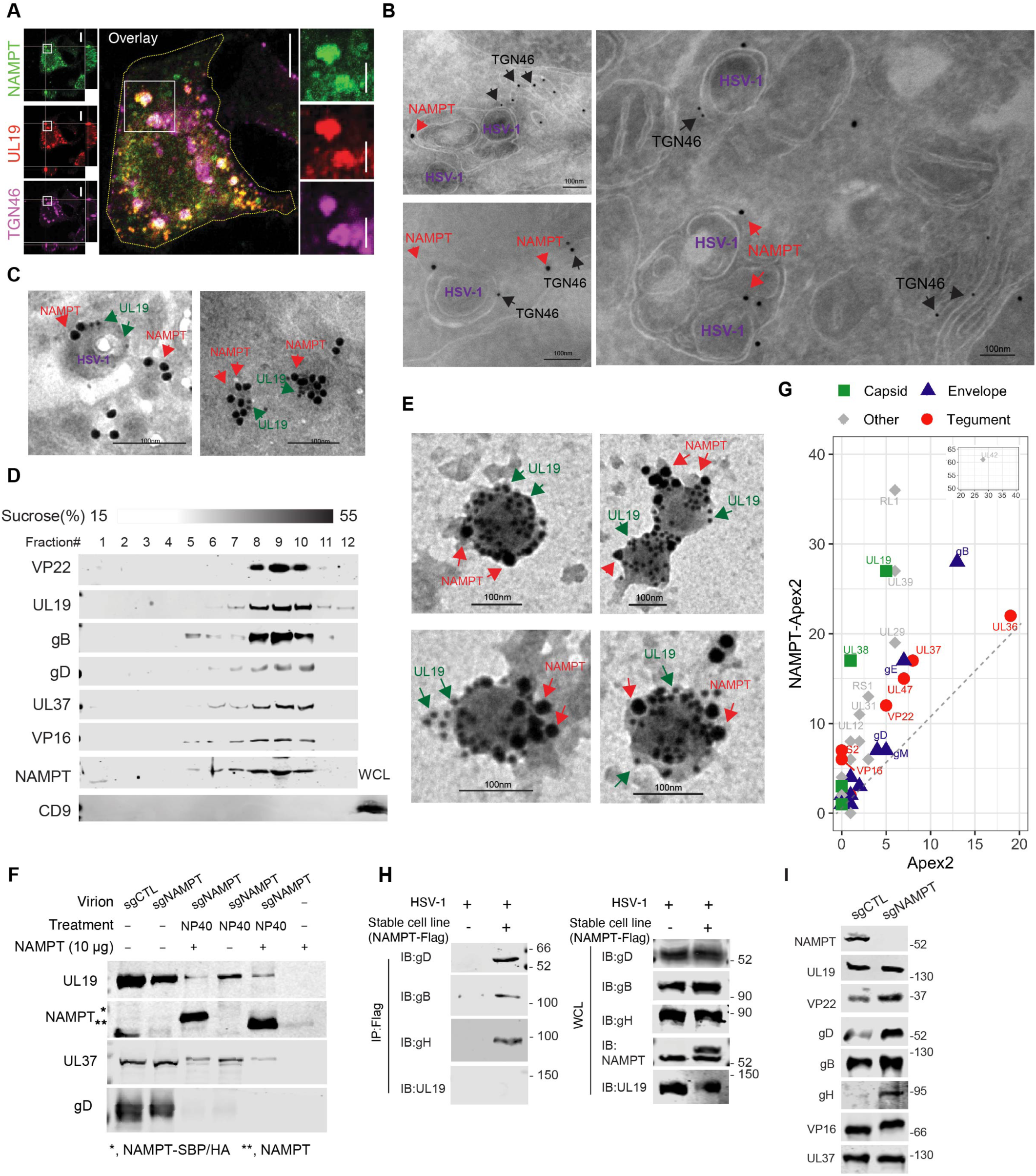
NAMPT is packaged in HSV-1 virions. (A to C) sgNAMPT HeLa cells reconstituted with HA-NAMPT were infected with HSV-1 (MOI=2) for 12 h. Immunofluorescence image of HSV-1-infected HeLa cells with antibodies to HA (NAMPT), UL19 and TGN46, with boxed region amplified and shown on the right (A). Immunogold staining followed by electron microscopy analysis with antibody against HA (NAMPT, 25 nm gold) and TGN46 (10 nm gold) (B) or HA (NAMPT, 25 nm gold) and UL19 (10 nm gold) (C). (D) Immunoblotting analysis of fractions of sucrose gradient ultracentrifugation with indicated antibodies. (E) Transmission electron microscopy analysis of purified HSV-1 virions after immunogold-staining with antibodies to UL19 (10 nm gold) and HA (NAMPT, 25 nm gold). (F) Interaction between NAMPT and de-enveloped HSV-1 virions analyzed by co-sedimentation in sucrose gradient ultracentrifugation and immunoblotting with indicated antibodies using pelleted virions. (G) Scatter plots of viral proteins identified by mass spectrometry after NAMPT-APEX2-mediated proximity ligation and normalization to the APEX2 control. (H) HSV-1 proteins interact with NAMPT by co-immunoprecipitation in HSV-1-infected 293T cells that stably express Flag-NAMPT. (I) Immunoblotting analysis of purified HSV-1 virions derived from sgCTL and sgNAMPT HepG2 cells infected with HSV-1 (MOI=0.1, 48 h.p.i.). Scale bar, 100 nm.

We then determined whether NAMPT is associated with HSV-1 virions. First, NAMPT was detected in fractions containing HSV-1 virions analyzed by sucrose gradient centrifugation and immunoblotting for viral structural components, including UL19, gB, gD, UL37, and VP16 (Figure 3D). These fractions were devoid of exosomes detectable with antibodies to well-established exosome markers, including CD9, CD81, Alix, and CD63 (Figure 3D and S4D). Using purified HSV-1 virions, we performed immunogold staining and found that NAMPT was abundantly detected in proximity to UL19, the major capsid protein of HSV-1 virions (Figure 3E and S4E). Interestingly, some virions showed little or no NAMPT staining, likely due to the incomplete permeabilization by NP40, and measured ∼150-200 nm in diameter, which is consistent with the size of enveloped HSV-1 virions.^21^ When treated with proteinase K, virion-associated NAMPT was resistant to proteinase K without detergent but became sensitive to proteinase K upon treatment with Triton X-100, demonstrating a sensitivity pattern similar to that of UL19 and UL37, which are capsid and tegument proteins packaged within the membrane, respectively (Figure S4F). To test whether NAMPT can associate with de-enveloped HSV-1 nucleocapsids, we performed a co-sedimentation experiment using sucrose ultracentrifugation as previously reported.^22^ We found that de-enveloped, but not intact HSV-1 virions, precipitated exogenous NAMPT purified from bacteria during ultracentrifugation (Figure 3F). These results show that NAMPT is packaged in the tegument compartment of HSV-1 virions.

To identify the viral proteins with which NAMPT interacts, we conducted ascorbic acid peroxidase (APEX)-mediated proximity labeling, biotin-based protein purification, and mass spectrometry analysis using HSV-1-infected HeLa cells expressing NAMPT-APEX2 (Figure S4G). Compared to APEX2, NAMPT-APEX2 expression resulted in the enrichment of several structural proteins, including tegument UL36, UL37, UL47, VP22, VP16, US2; capsid UL19 and UL38; and envelope gB, gE, gD, and gM glycoproteins (Figure 3G). When co-immunoprecipitation was performed to confirm these interactions, we found that NAMPT interacts with several glycoproteins, including gB, gD, and gH in HSV-1-infected HeLa cells; and with tegument proteins, including UL21, UL37, UL47, VP16, and VP22 in transfected 293T cells (Figure 3H, S4H, and S4I). No interaction between NAMPT and UL19, UL38, or UL18 was detected under similar conditions (Figure 3H and S4J). We then purified HSV-1 virions derived from wildtype and NAMPT-knockout HepG2 cells and analyzed them by immunoblotting. This showed that, when HSV-1 virions were normalized with the UL19 major capsid protein, higher levels of VP22, gD, gH, and to a lesser extent, gB, were detected in virions produced from NAMPT-knockout HepG2 cells compared to those produced from wildtype HepG2 cells (Figure 3I). No significant change was observed for UL37. Notably, the migration of VP16 was clearly retarded by loss of NAMPT, suggesting possible post-translational modifications. Increases in VP22, gD, and gB in virions produced from NAMPT-knockdown HepG2 cells were also observed (Figure S4K). These results demonstrate that NAMPT restricts the incorporation of tegument proteins and glycoproteins into HSV-1 virions.

### NAMPT is a protein phosphoribosyl-hydrolase (phosphoribosylase)

Given that NAMPT interacts with multiple proteins and alters their incorporation into HSV-1 virions, we reasoned that this may depend on its activity to remove phosphoribosyl moieties. Inspired by previous studies demonstrating that other metabolic enzymes act on proteins post-translationally,^23,24^ we hypothesized that NAMPT catalyzes the hydrolysis of phosphoribosyl moieties of viral and cellular proteins to restrict HSV-1 replication. To differentiate it from the phosphatase activity toward the phosphoribosyl moiety, we designate the glycohydrolase activity of NAMPT as a phosphoribosylase. To test this hypothesis, we reconstituted NAMPT-knockout HeLa cells with wildtype NAMPT or the NAMPT-H247E and -D219A mutants that are impaired for phosphoribosyl transfer and substrate (NAM)-binding, respectively, for HSV-1 lytic replication.^25^ Notably, although the NAMPT-H247E and -D219A mutants are functionally compromised, significant residual enzyme activity could be detected by an *in vitro* NAD^+^ synthesis assay (Figure S5A). Interestingly, wildtype NAMPT, but not the NAMPT-H247E mutant, inhibited HSV-1 replication (Figure 4A and S5B). The NAMPT-D219A mutant, however, more robustly reduced HSV-1 replication than wildtype NAMPT, despite that all three were being expressed at similar levels. Thus, the enzyme activity required for NAMPT-mediated restriction of HSV-1 is distinct from its activity in the salvage NAD^+^ synthesis. Next, we performed a modified phosphoproteomics protocol, consisting of phosphopeptide enrichment and LC-MS analysis to identify phosphoribosylated peptides using HSV-1-infected HeLa cells expressing the NAMPT-H247E mutant (Figure 4B). Compared to control HeLa cells, the total abundance of phosphoribosylated peptides roughly doubled in HeLa cells with NAMPT-H247E expression, as indicated by the total abundance and percentage of phosphoribosylated peptides (Figure S5C). This analysis uncovered a panel of phosphoribosylated peptides of viral proteins, including glycoproteins gB, gD, and gH; tegument proteins VP22, VP16, UL21, UL37, UL46, and UL47; and capsid proteins UL19, UL17, and UL18 (Figure S5D). Interestingly, three ADP-ribosylated peptides of VP22 were identified as well. All ribosylated sites contained either phosphoribose or ADP-ribose, but not both, suggesting that phosphoribose and ADP-ribose mark distinct biochemical and functional entities of these modified peptides in HSV-1 infection. Notably, these phosphoribosylated proteins largely overlap with those NAMPT-binding partners identified by APEX2-mediated proximity labeling (Figure 3G). Among these phosphoribosylation sites, most are aspartates or glutamates, and a few are arginines, which is consistent with the profile of ADP-/phospho-ribosylation sites that have been previously reported (Figure 4C).^26–28^ Phosphoribosylation of these viral peptides was validated by parsing the mass spectrum of each peptide (Figure 4D). Phosphoribosylated peptides of cellular proteins were identified and their roles in HSV-1 replication were not investigated in this study.

**Figure 4.**
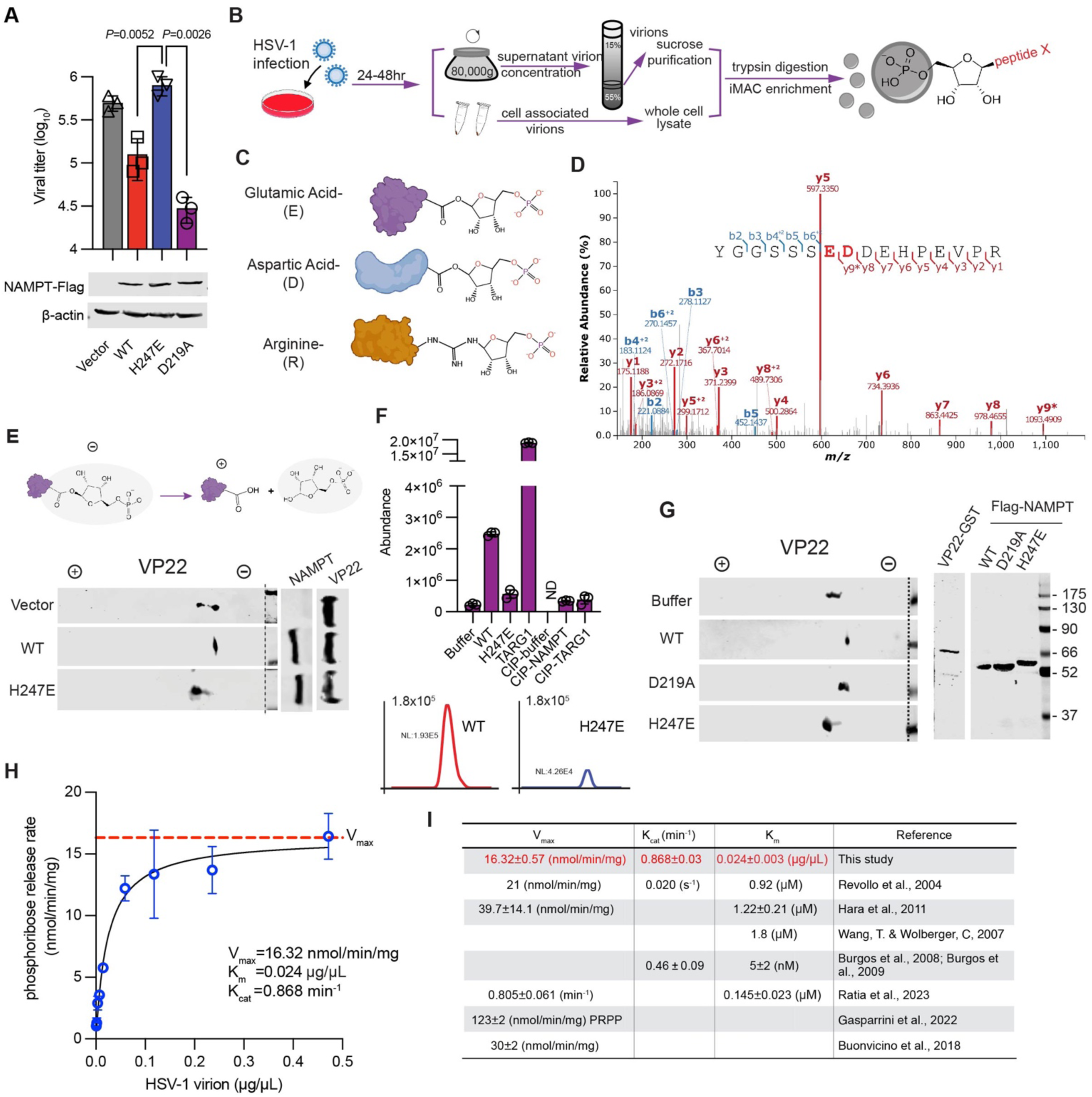
NAMPT is a protein phosphoribosylase. (A) HSV-1 titer in the medium of sgNAMPT HeLa cells reconstituted with wildtype, H247E and D219A mutants of NAMPT at 24 h.p.i. (MOI=0.1), while whole cell lysates were analyzed for the expression of NAMPT proteins (bottom panels) by immunoblotting with indicated antibodies. (B) Schematic illustration of the procedure to identify phosphoribosylated peptides. (C) Schematic illustration of phosphoribose linked to glutamate, aspartate and arginine residues of peptides. (D) Tandem mass spectrum of the VP22 peptide containing phosphoribosylated E74 and D75 that are highlighted in red. (E) Two-dimensional gel electrophoresis and immunoblotting analysis of 293T cells expressing GST-VP22 with wildtype (WT) or the H247E mutant of NAMPT. (F-H) Elution profile by liquid chromatography (F) of phosphoribose cleaved by NAMPT, and two-dimensional gel electrophoresis and immunoblotting analysis of GST-VP22 (G) of the in vitro phosphoribosylase reaction, and the phosphoribosylase activity of NAMPT analyzed by released phosphoribose that is quantitatively determined by liquid chromatography-mass spectrometry (H). (I) The *V_max_*, *K_cat_* and *K_m_* of NAMPT phopshoribosylase activity determined in this study are compared to its phosphoribosyltransferase activity (in NMN synthesis) defined by previous studies. Statistical significance was calculated using unpaired two-tailed Student’s *t*-tests.

VP22, one of the most abundant tegument proteins, is highly phosphoribosylated as indicated by the multiple peptides carrying phosphoribose. We then analyzed the charge status of VP22 by two-dimensional gel electrophoresis with the overexpression of wildtype NAMPT or the NAMPT-H247E mutant. We found that wildtype NAMPT diminished the more negatively charged species, while the NAMPT-H247E mutant abolished the more positively charged species, suggesting that NAMPT-H247E functions as a dominant negative mutant to inhibit the phosphoribosylase activity of endogenous NAMPT (Figure 4E). Conversely, re-expression of NAMPT in NAMPT-knockout 293T cells shifted VP22 toward the negative pole of the gel strip, providing support for NAMPT’s phosphoribosylase activity against VP22 (Figure S5E). Furthermore, purified wildtype NAMPT, but not the NAMPT-H247E mutant, released phosphoribose in an *in vitro* assay (Figure 4F, S5F, and S5G). Compared to NAMPT, the terminal ADP-ribose glycohydrolase 1 (TARG1) demonstrated higher activity, which is likely due to its lower substrate selectivity. When VP22 was analyzed by two-dimensional gel electrophoresis, we found that wildtype NAMPT and the NAMPT-D219A mutant, but not the NAMPT-H247E mutant, shifted VP22 toward the negative end of gel strip, consistent with its phosphoribosylase activity against HSV-1 proteins (Figure 4G). With VP22 defined as a substrate of NAMPT, we generated the VP22-E257A mutant and found that it was not shifted by the NAMPT-H247E mutant as analyzed by two-dimensional gel electrophoresis, further supporting that NAMPT acts as a phophoribosylase for VP22 (Figure S5H). To further characterize the protein phosphoribosylase activity of NAMPT, we performed an *in vitro* assay using purified HSV-1 virions, because purified GST-VP22 was not sufficient to achieve maximal velocity (V_max_) of the phosphoribosylase activity of NAMPT. When released phosphoribose was quantitatively determined by LC-MS, NAMPT demonstrated a V_max_ of 16.32 nmole/min/mg and a K_cat_ of 0.864 min^-1^ toward HSV-1 virions (Figure 4H, 4I, and S5I). The V_max_ of phosphoribosylase activity of NAMPT is very close to that of its phosphoribosyltransferase activity that was previously reported.^29^ However, the K_cat_ of the phosphoribosylase activity is much higher than that of its phosphoribosyltransferase activity, presumably because phosphoribosyl hydrolysis is energy-favored (Figure 4I). These results collectively support the conclusion that NAMPT is a protein phosphoribosylase.

### Phosphoribose of structural proteins promotes their incorporation into HSV-1 virions

Having defined NAMPT as a protein phosphoribosylase and identified a panel of phosphoribosylated viral proteins, we then assessed the charge status of HSV-1 proteins by expressing the dominant-negative NAMPT-H247E mutant. This assay showed that NAMPT-H247E expression reduced the charge of several HSV-1 proteins, including gB, gD, gH, and UL21, while it had no significant effect on that of UL17 and UL38 (Figure S6A). A minor increase in charge was observed for VP16, UL18, UL19, UL37, and UL47 under similar conditions (Figure S6A). These results suggest that NAMPT potentially acts as a phosphoribosylase for multiple viral proteins.

To determine whether phosphoribosyl moieties affect viral protein incorporation, we first analyzed the charge status of gD expressed in HSV-1-infected cells and of HSV-1 virions. A total of six species with distinct charge statuses were detected for gD expressed in HSV-1-infected cells and extracellular HSV-1 virions. Interestingly, gD packaged in HSV-1 virions exhibited more negatively charged species than that expressed in HSV-1-infected cells (Figure S6B and S6C). This was reflected by the reduced abundance of the most positively charged species and a corresponding increase in the more negatively charged species of the virion-associated gD. We then analyzed the effect of NAMPT knockout on the charge status of gD, and two-dimensional gel electrophoresis revealed a total of 10 species of gD by charge status in HSV-1-infected cells and purified virions (Figure 5A). In control HepG2 cells, gD packaged in HSV-1 virions was more negatively charged than that expressed in HSV-1-infected cells. Remarkably, NAMPT knockout in HepG2 cells resulted in a major shift toward gD species that were highly negatively charged in both HSV-1-infected cells and HSV-1 virions. Furthermore, the more negatively charged species were enriched in HSV-1 virions compared to those expressed in HSV-1-infected cells. Quantification of the distinct gD species showed that loss of NAMPT had a significant impact on the charge status of gD, resulting in the packaging of more negatively charged species in HSV-1 virions (Figure 5B). These results support the conclusion that gD is likely phosphoribosylated, and that highly phosphoribosylated gD species are incorporated into HSV-1 virions.

**Figure 5.**
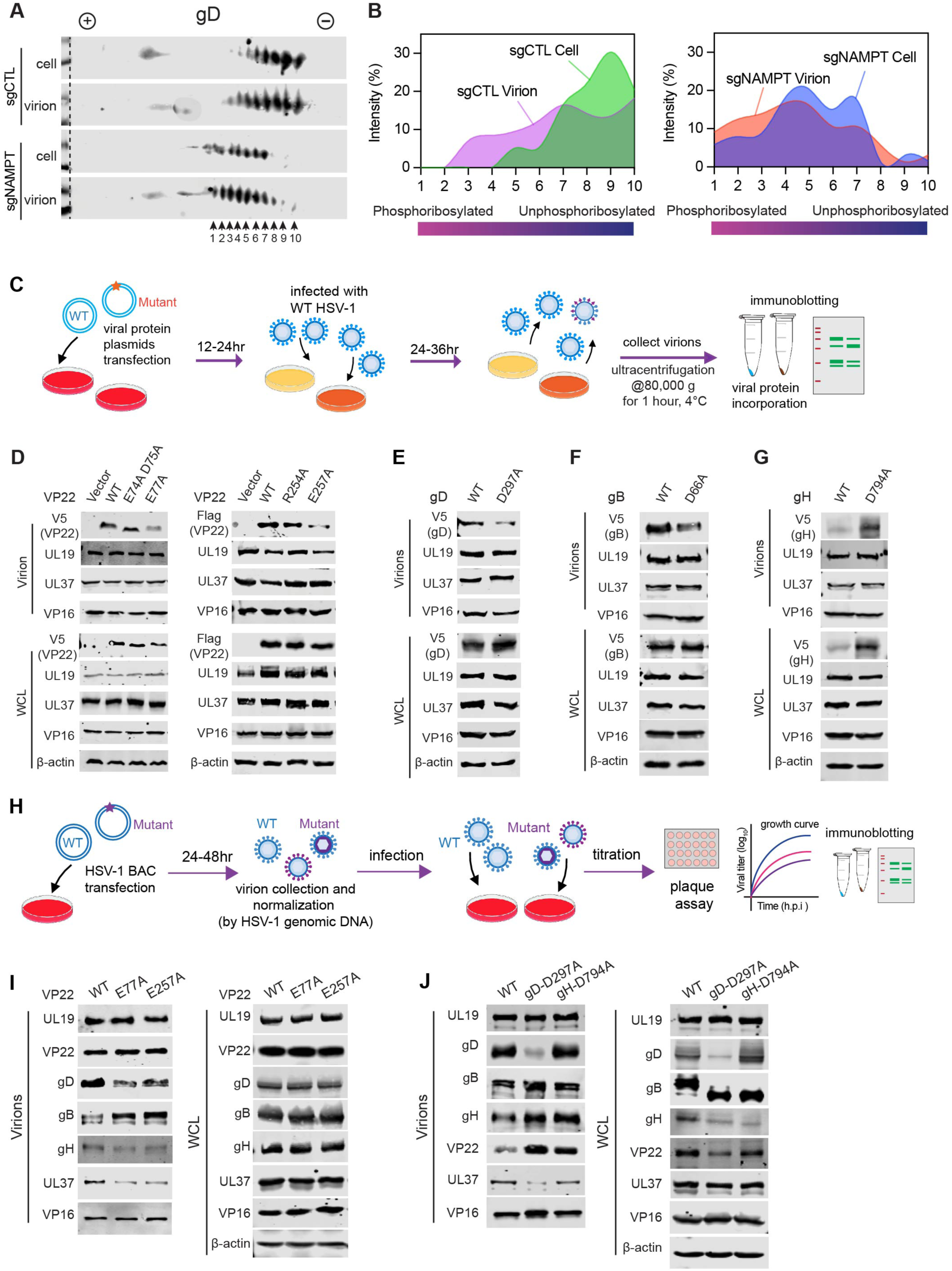
Phosphoribose of structural proteins promotes their virion incorporation. (A) Two-dimensional gel electrophoresis and immunoblotting analysis of purified HSV-1 virions with antibody to gD (left), and quantification by densitometry of the gD species with distinct charge as indicated by arrows in (right). (B) Schematic illustration of the procedure to evaluate the virion incorporation of HSV-1 mutant proteins. 293T cells transiently expressing HSV-1 proteins (wildtype and phosphoribosylation-resistant mutants) were infected with HSV-1 (MOI=0.2) and extracellular virions were harvested at 36 h post-infection for immunoblotting analysis to examine HSV-1 structural proteins in reference to their expression in 293T cells. (C to F) Wild-type and phosphoribosylation-resistant mutants of HSV-1 proteins, including VP22 (C), gD (D), gB (E) and gH (F), in HSV-1 virions and expressed in HSV-1-infected cells were analyzed by immunoblotting with indicated antibodies as indicated. All HSV-1 proteins were tagged with the V5 epitope. (G) Schematic illustration of engineering and characterizing recombinant HSV-1 using BAC. (H and I) Immunoblotting analysis of purified virions of recombinant HSV-1 containing VP22-E77A or - E257A (H), gD-D297A or gH-D794A (I) compared to wildtype HSV-1, with HSV-1 virion normalized to UL19, and whole cell lysates of infected HeLa cells as controls with indicated antibodies.

To further profile the effect of phosphoribosylation-resistant mutations on HSV-1 proteins, we generated mutant constructs of structural proteins and analyzed their incorporation into HSV-1 virions. To do that, transfected 293T cells were infected with wildtype HSV-1 to obtain chimeric virions that carry both wildtype and a mutant form of the structural protein in the medium (Figure 5C). Virion incorporation of the mutant protein was analyzed by immunoblotting. When wildtype VP22 and its mutants incorporated in extracellular HSV-1 virions were normalized with the UL19 major capsid protein, we found that mutations of E77A and E257A reduced VP22 virion incorporation without affecting VP22 expression (Figure 5D). By contrast, phosphoribosylation-resistant mutations E74A, D75A, and R254A had no effect on VP22’s virion incorporation, indicating the site-specific effect of phosphoribosylation. Phosphoribosylation-resistant mutations of gB and gD reduced their incorporation in HSV-1 virions, while the mutation of gH increased its expression in transfected/infected cells and subsequent virion incorporation (Figure 5E-5G). Mutations of UL37 (E238A), VP16 (E6A, D7A, D11A, and E314A), UL18 (D9A and D45A), and UL47 (E636A and D669A) had no significant effect on their virion incorporation (Figure S6D-S6G). Interestingly, UL32-E353A did not affect its own expression in cells or incorporation into HSV-1 virions, although it impeded the virion incorporation of UL37, but not VP16 (Figure S6H). This effect may stem from the slightly lower UL37 expression in HSV-1-infected cells. These results show that phosphoribosylation of VP22 and glycoproteins promotes their incorporation into HSV-1 virions.

Next, we engineered recombinant HSV-1 carrying mutations abolishing phosphoribosylation via bacteria artificial chromosome (BAC) and examined the virion incorporation of the structural proteins (Figure 5H). In parallel, we also generated recombinant HSV-1 that contained mutations abolishing the ADP-ribosylation of VP22, i.e., R86A, E149A, and E234A (Figure S5D). All mutations were validated by sequencing of the targeted region of the HSV-1 genome. All BAC DNAs, except that carrying gB-D66A, yielded infectious recombinant HSV-1 when transfected into Vero cells. With recombinant HSV-1, we analyzed the virion incorporation of HSV-1 structural proteins by immunoblotting. As shown in Figure 5I, E77A and E257A had no effect on the virion incorporation of VP22 and VP16 but reduced that of gD, gH, and UL37, while increasing the more slowly migrating (presumably more glycosylated) species of gB, despite all viral proteins being equally expressed in HSV-1-infected cells. In cells infected with HSV-1 containing gD-D297A, gD expression was greatly reduced while gB migrated much faster than that produced from the HSV-1 wildtype, likely reflecting its lack of proper glycosylation (Figure 5J). For HSV-1 virions containing gD-D297A, gD-D297A and UL37 were greatly reduced compared to wildtype HSV-1, while virion incorporation of gH and VP22, and to a lesser extent, that of gB and VP16, clearly increased. Importantly, both gD-D297A and gH-D794A altered the expression of glycoproteins and tegument proteins in HSV-1-infected cells, which do not correlate with their virion incorporation. Though gH-D794A had the same effect on gB expression in HSV-1-infected cells as gD-D297A, it demonstrated a distinct impact on the virion incorporation of gH and VP22 (Figure 5J). Thus, phosphoribosylation of VP22 and these glycoproteins affects the processing (e.g., glycosylation) and virion incorporation of glycoproteins.

### Phosphoribose of structural proteins enhances HSV-1 entry

Glycoproteins, including gB, gD, gH, and gL, constitute a multi-component machinery that mediates HSV-1 entry.^30,31^ Along with the observation that mutations in VP22 affect the virion incorporation of multiple glycoproteins, phosphoribosylation of three out of four key entry glycoproteins prompted us to examine its role in HSV-1 entry. We first generated chimeric HSV-1 carrying both wildtype and phosphoribosylation-resistant mutants of gB, gD, and gH for the entry assay. 293T cells over-expressing wildtype and phosphoribosylation-resistant mutants of gB, gD, and gH were infected with wildtype HSV-1 to produce chimeric HSV-1 that carries both wildtype and mutant glycoproteins (Figure 6A). When HSV-1 entry was analyzed by intracellular viral genomes, we found that phosphoribosylation-resistant gD and gB reduced HSV-1 entry by ∼85% and 60%, respectively, while phosphoribosylation-resistant gH had no significant effect (Figure 6B). Given the reduced virion incorporation of gD-D297A and gB-D66A (Figure 5C and 5D), the effect of these phosphoribosylation-resistant mutants on HSV-1 entry is likely under-estimated. Next, we used recombinant HSV-1 carrying phosphoribosylation-resistant mutants of gD and gH for the entry assay, although the insufficiency of HSV-1 containing gB-D66A precluded a quantitative analysis of its effect on HSV-1 entry (Figure S5J). Indeed, recombinant HSV-1 containing gD-D297A demonstrated a 50-75% reduction in entry, while the gH-D794A mutant showed no significant deficiency compared to wildtype HSV-1 (Figure 6C and 6D). Similarly, recombinant HSV-1 carrying phosphoribosylation-resistant mutations, i.e., E77A or E257A, in VP22 also demonstrated reduced HSV-1 entry compared to wildtype HSV-1 (Figure 6E).

**Figure 6.**
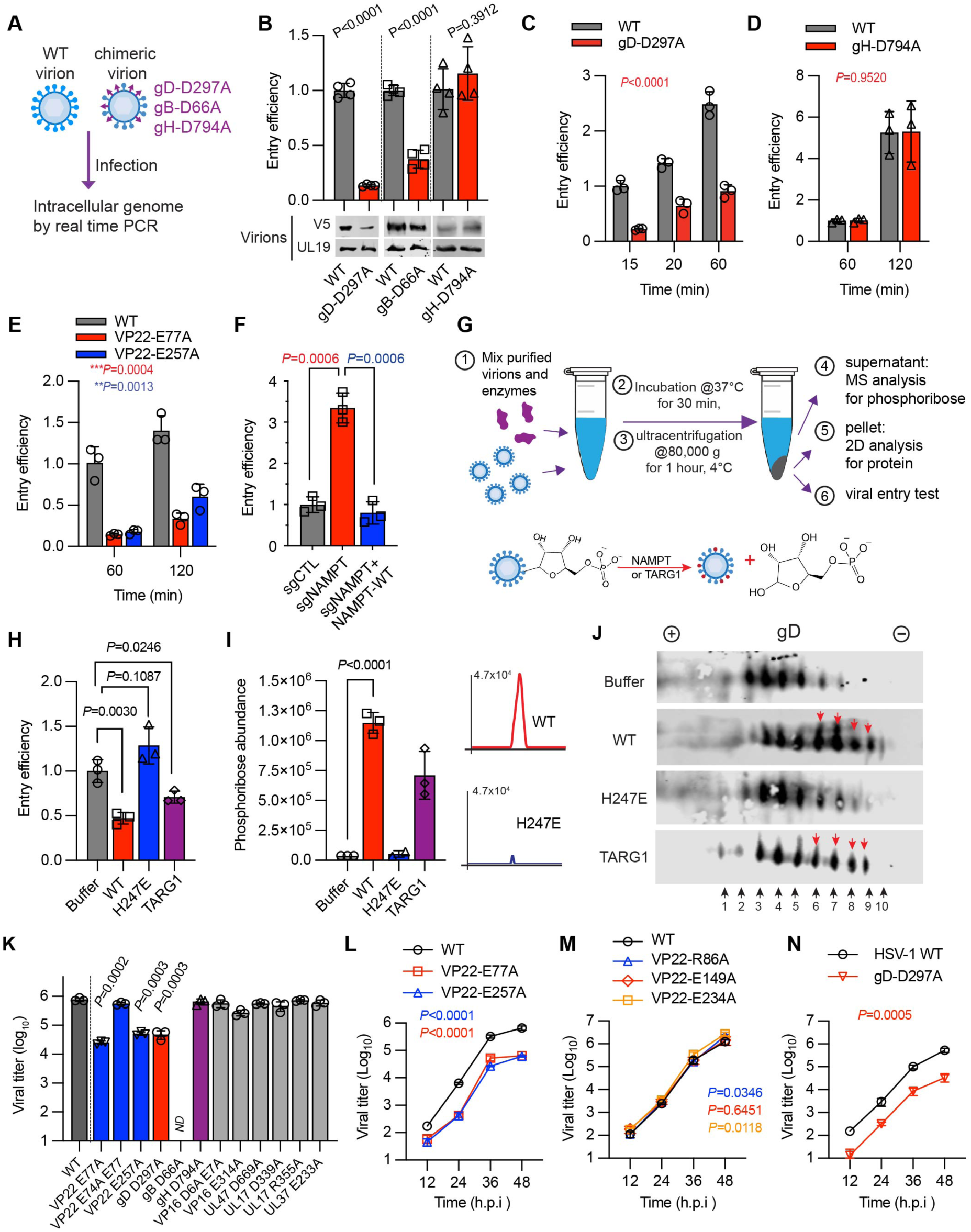
Phosphoribose of structural proteins in HSV-1 entry and replication. (A) Schematic illustration of entry analysis using WT and chimeric HSV-1 virions containing gD-D297A, gB-D66A or gH-D794A. (B) Entry analysis of HSV-1 using HeLa cells as target cells, with HSV-1 produced from 293T cells transiently expressing wildtype gB, gD and gH or their corresponding phosphoribosylation-resistant mutants as indicated. (C to E) Entry analysis of wildtype (WT) recombinant HSV-1 and that containing VP22-E77A or -E257A mutations (C), gD-D297A (D) or gH-D794A (E). (F) Entry analysis of HSV-1 produced from sgCTL, sgNAMPT HeLa cells, or sgNAMPT HeLa cells reconstituted with NAMPT. (G to J) Schematic illustration of the in vitro phosphoribosylase reaction and subsequent functional or biochemical analysis using purified HSV-1 virions (G). Entry analysis (H), quantification of phosphoribose released in the phosphoribosylase reaction by mass spectrometry (I), and two-dimensional gel electrophoresis and immunoblotting analysis with anti-gD antibody (J) of HSV-1 virions treated with NAMPT, NAMPT-H247E or TARG1. (K) Titer of recombinant HSV-1 carrying indicated mutations in the medium of HepG2 cells at 48 h.p.i. (MOI=0.1). Note, recombinant HSV-1 carrying gB-D66A failed to amplify. (L to N) Growth curve of wildtype HSV-1 and that containing VP22 E77A or -E257A (phosphoribosylation-resistant, L), R86A, E149A, E234A (ADP-ribosylation-resistant, M) and that containing gD-D297A (N) in the medium of HepG2 cells at MOI=0.1, was determined by plaque assay using Vero cells. Statistical significance was calculated using unpaired two-tailed Student’s *t*-tests, for B, F, H, I and K; two-way ANOVA for C, D, E, L, M and N.

To biochemically demonstrate the role of phosphoribosyl moieties on glycoproteins, we first compared HSV-1 virions produced from NAMPT-knockout HepG2 cells to those produced from wildtype HepG2 cells by the entry assay. We found that HSV-1 virions produced from NAMPT-KO cells demonstrated ∼2.5-fold more entry than those produced from wildtype HeLa cells, and the re-expression of NAMPT in NAMPT-knockout HeLa cells completely abolished the increase in HSV-1 entry induced by NAMPT knockout (Figure 6F). Next, we treated HSV-1 virions produced from NAMPT-knockout HepG2 cells with either NAMPT or the NAMPT-H247E mutant. We examined HSV-1 entry, quantified phosphoribose, and analyzed glycoproteins by two-dimensional gel electrophoresis (Figure 6G). This analysis showed that wildtype NAMPT and TARG1, but not NAMPT-H247E, reduced HSV-1 entry by ∼55% (Figure 6H). Interestingly, the NAMPT-mediated inhibition requires ATP, which is consistent with the ATP-dependent activation of NAMPT (Figure S7A). Consistent with this result, NAMPT, but not the NAMPT-H247E mutant, released a significant amount of phosphoribose from HSV-1 virions, as quantified by LC-MS (Figure 6H). Furthermore, NAMPT and TARG1 also increased the relative amount of the more positively charged species of gD and gB in HSV-1 virions, as shown by two-dimensional gel electrophoresis, but the NAMPT-H247E mutant had little or no effect, supporting the phosphoribosylase activity of NAMPT toward HSV-1 glycoproteins (Figure 6I and S7B). When NAM was added to assess the phosphoribosyltransferase activity of NAMPT, we found that NAMPT extracted phosphoribose, but failed to synthesize NMN (Figure S7C). As expected, NAMPT catalyzed the NAM to NMN conversion when exogenous PRPP was added. Thus, this result also suggests that the released phosphoribose was not used to synthesize NMN or PRPP in the phosphoribosylase reaction. These findings show that phosphoribose of glycoproteins and VP22 promotes HSV-1 entry.

### Phosphoribose of structural proteins facilitates HSV-1 lytic replication

Our studies involving chimeric and recombinant HSV-1 show that phosphoribose is important for the virion incorporation of structural proteins and subsequent entry. Thus, we examined the effect of phosphoribose on HSV-1 lytic replication using recombinant HSV-1 carrying phosphoribosylation-resistant mutations. Multi-step growth curve showed that phosphoribosylation-resistant mutations in VP22, gB, gD, and VP16 significantly reduced HSV-1 replication, whereas mutations in gH, UL17, UL37, and UL47 had no apparent effect on HSV-1 replication, as measured by extracellular viral titer (Figure 6J-L, S7D, and S7E). Notably, mutations in VP22 (E77A and E257A) and in gD reduced extracellular HSV-1 titer by ∼10-50-fold (Figure 6J). The gB-D66A mutation severely impaired HSV-1 lytic replication to the extent that amplification of the recombinant HSV-1 was unsuccessful. Conversely, an HSV-1 revertant containing gB-A66D behaved similarly to wildtype HSV-1, excluding the contribution of undesired mutations (Figure S7F). By contrast, the mutations abolishing the three ADP/polyADP-ribosylations of VP22 had no detectable effect on HSV-1 lytic replication (Figure 6J). These results collectively support a distinct role of phosphoribosyl modification in HSV-1 lytic replication. Together, these results characterized a number of phosphoribosylation-resistant mutations of tegument and glycosylated proteins in HSV-1 virion assembly and entry (Figure S7G), demonstrating pivotal roles of phosphoribosylation in HSV-1 infection.

## DISCUSSION

Viral infections represent major challenges to public health, as evidenced by the recent COVID pandemic, and the development of innovative therapeutic strategies will require a better understanding of host antiviral defense. Known as a metabolic phosphoribosyltransferase in the salvage NAD^+^ synthesis, NAMPT inhibits a handful of enveloped viruses, including West Nile virus, Venezuelan equine encephalitis virus, and human immunodeficiency virus (HIV) 1.^18,32^ In this study, we show that NAMPT acts as a phosphoribosylase toward several viral structural proteins to restrict HSV-1 lytic replication, ascribing unprecedented enzymatic activity to a key metabolic enzyme. Previously, antiviral metabolic enzymes were reported to impede viral replication processes by modifying key metabolites, such as SAMHD1 for HIV and viperin for various RNA viruses,^4,5^ or by evading the host immune response, as observed with glutamine aminotransferases.^33,34^ NAMPT recycles NAM to regenerate NAD^+^ that is crucial for further downstream biological processes, such as DNA damage response, epigenetic regulation, and gene expression, all of which likely contribute to an efficient productive infection of HSV-1. By contrast, NAMPT de-phosphoribosylates viral, and likely cellular proteins to hinder the assembly and subsequent infection of HSV-1 virions, thus qualifying as a “moonlighting” enzyme.^35^ The distinct activities of NAMPT in salvage NAD^+^ synthesis and protein dephosphoribosylation is substantiated by the observation that the NAMPT-D219A mutant demonstrated residual activity in salvaging NAM, while higher activity to restrict HSV-1 lytic replication. Interestingly, ADP-ribose metabolic enzymes such as TARG1 were originally discovered to act on proteins as a terminal glycohydrolase and recently found to have demonstrated similar activity towards RNA and DNA.^36,37^ The phosphoribosyltransferase and phosphoribosylase activities of NAMPT toward small molecules and proteins, respectively, may also play a role in other microbial infections and human diseases.^8,17^ This begs for further investigation into the dynamic regulation of these two activities.

NAMPT catalyzes the synthesis of NAD^+^ via salvaging NAM, which is the byproduct of predominant NAD^+^-consuming enzymes such as PARPs, SIRTs, and SARM1. Considering the essential roles of NAD^+^ in diverse biological processes, it is surprising that NAMPT deficiency increased HSV-1 lytic replication. Indeed, addition of NR to the medium effectively restores intracellular NAD^+^ concentration and further boosts HSV-1 replication in NAMPT-knockout or -depleted cells, suggesting that the antiviral activity of NAMPT is likely underestimated. Previous studies showed that global deletion of NAMPT is lethal in mice, largely due to deficits in neurodevelopment.^38,39^ In a global inducible knockout mouse strain, significant depletion of NAMPT was observed only in the liver and spleen, but not in the heart, lung, and kidney upon induction with tamoxifen. This may explain the lack of apparent abnormalities in these mice. NAMPT deletion in the liver reduced hepatic abundance of NAD^+^ by ∼50%, which was restored by exogenous NR through intraperitoneal injection. This scenario resembles NAMPT-depleted cells cultured with the medium containing regular FBS. Consistently, NR administration to NAMPT-knockout mice also increased HSV-1 replication in the NAMPT-depleted liver. Given the short duration of HSV-1 lytic replication in mice, we profiled innate immune gene expression in cells and the liver with NAMPT depletion. We found that NAMPT depletion slightly increased the expression of some genes, possibly due to elevated HSV-1 lytic replication. These results support the notion that the loss of NAMPT does not grossly impair innate immune defense against HSV-1 in the NAMPT-KO mice, and that NR supplementation further increases HSV-1 lytic replication in NAMPT-depleted tissues.

In metazoans, phosphoribose on proteins was previously identified through in-depth analysis of phosphoproteomic datasets, but no related functions have been described as of yet.^27^ SdeA, an effector protein secreted by *Legionella pneumophila*, catalyzes the phosphoribosylation of ubiquitin that is subsequently transferred to a target protein, thereby impairing the cellular ubiquitination cascade and related biological processes such as TNF-α signaling.^40^ Considered as relics of ADP/poly(ADP)- ribosylation, phosphoribose moieties are likely derived from ADP/polyADP-ribose through enzymes like NUDT16, which has demonstrated the ability to convert ADP/polyADP-ribose into phosphoribose in assembled reactions.^14^ However, such an activity of NUDT16 and enzymes alike needs to be confirmed in cells and the biological significance of this enzymatic activity has not been reported. Among the three ADP-ribosylation sites and four phosphoribosylation sites identified within VP22, none of these sites contain both phosphoribosyl and ADP-ribosyl moieties. This finding supports the notion that ADP-ribosyl and phosphoribosyl units represent distinct biochemical and functional entities of VP22. Indeed, mutational analyses further showed that two phosphoribosylation-resistant mutations, but none of the three ADP-ribosylation-resistant mutations, severely impaired HSV-1 lytic replication. Furthermore, the phosphoribosylation of the three out of four glycoproteins of the primary entry complex and the VP22 tegument protein functionally converge to impinge on virion assembly and the entry of new infection, signifying the pivotal roles of phosphoribosylation in late stages of HSV-1 lytic replication. This conclusion is substantiated by electron microscopy analysis showing that NAMPT acts in the TGN during HSV-1 tegumentation. Notably, the biology of protein phosphoribosylation is poorly understood. Herpesvirus infection therefore provides an excellent model system to address key questions concerning protein phosphoribosylation in assembly and egress in HSV-1 infection in particular and the fundamental biological processes in metazoans in general.

## Acknowledgements

Acknowledgements. We thank Dr. Shin-Ichiro Imai (Washington University) for providing the *Nampt^fl/fl^*mice, Dr. Jon Hao (Poochon Scientific) for proteomics analysis, Dr. Wandy Beatty (Washington University) for electron microscopy analysis, Drs. Weiming Yuan (University of Southern California), David Knipe (Harvard Medical School) and Richard Longnecker (Northwestern University) for providing HSV-1 plasmids, Dr. Jun Zhao (Cleveland Clinic Foundation) for guidance on generating HSV-1 mutant viruses, Dr. Gary Cohen (University of Pennsylvania) for anti-gH antibody, Dr. Chunfu Zhen (University of Calgary) for anti-VP22 antibody, and Mr. Brett Lomenick (CalTech) for processing MS samples. We are grateful to Yuzheng Zhou, Stephanie Rice, Drs. Jessica Carriere and Jun Xiao for their assistance. This work was partly supported by a startup fund from the Herman Ostrow School of Dentistry of USC and grants from NIH (AG070904, CA285192 and AI180537) and Infectious Disease Society of America Foundation (Microbial Pathogenesis in AD).

## Author contributions

Conceptualization: PF, SZ, QC; Methodology: SZ, NX, YL, QC, TW, PF, TC; Investigation: SZ, NX, YL, QC, ACS, TW, YR, SL, AS; Funding acquisition: PF, CB; Project administration: PF, CH; Supervision: PF, CB, CH, TC; Writing – original draft: PF, SZ; Writing – review & editing: PF, SZ, CB, CQ.

## CONTACT FOR REAGENT AND RESOURCE SHARING

Further information and requests for resources and reagents should be directed to and will be fulfilled by the Lead Contact, Pinghui Feng.

## Methods and Materials

### Cell Culture

HEK293T, HepG2, HeLa, Vero, MEF and HFF cells were purchased from ATCC and cultured in DMEM (HyClone) supplemented with 10% heat-inactivated FBS (Gibco), 1% penicillin/streptomycin (Corning) and maintained in the humidified incubator with 5% CO2 at 37°C. NAMPT knockout and control cells, sgNAMPT and sgCTL, were generated using CRISPR-Cas9 system. In brief, sgNAMPT or sgCTL constructs (pL-CRISPR-EFS-PAC, Addgene) and lentiviral packaging plasmids pMD2.g and psPAX (Primers and Constructs were described in Table S2) were transfected into 293T cells for lentivirus production. At 48 hours post transfection, the medium containing sgNAMPT or sgCTL lentivirus was collected, filtered through 0.22 μm membrane, and added to 293T, HepG2, HeLa cells for spinfection with 8 μg/mL polybrene at 1,000 *g* for 45 min. At 48 hours post-infection, puromycin was used to select cells at the final concentration of 2 μg/mL. To select single colony, pooled puromycin resistant cells were diluted into 96 well plates to achieve one cell per well. Each colony was expanded and further validated via immunoblotting and genomic sequencing. The sgNAMPT cells were maintained in the complete DMEM medium supplemented with 0.1 mM nicotinamide riboside (NR). NAMPT is critical for NAD^+^ regeneration and cell survival. The proliferation of NAMPT-depleted cells is likely due to NR in the fetal bovine serum (FBS). Indeed, when NAMPT-depleted HeLa cells were grown in medium containing dialyzed FBS, NAD^+^ levels decreased gradually and continuous culture with dialyzed FBS induced cell death (data not shown). By contrast, sgNAMPT HeLa cells continued to grow in medium with normal FBS.

To generate the reconstituted cell lines, sgNAMPT cells were spinfected with lentiviruses carrying plasmids: NAMPT-Flag, NAMPT-V5, NAMPT-HA, NAMPT-H247E-Flag or NAMPT-D219A-Flag. Expression of NAMPT and mutants were validated via immunoblotting with antibodies to NAMPT and designated tags.

### Viruses

Human herpes simplex virus 1 (strain 17) (HSV-1) was amplified in Vero cells. To perform HSV-1 infection and determine virus growth curve, control and NAMPT-knockdown or -knockout cells were infected with HSV-1 at MOI=0.1 in FBS-free DMEM medium for 2 hours. Fresh DMEM medium with 10% dialyzed FBS, supplemented with 0.1 mM NR, was used to replace the infection medium. The medium and cells were collected separately at 12, 24, 36, 48 and 72 hours post infection for plaque assay to determine viral titer. HSV-1 titer was determined by plaque assay on Vero monolayer. Briefly, 10-fold serial-diluted virus-containing medium (FBS-free DMEM) was added to Vero cells. After 2 hours of incubation in 5% CO2 at 37 °C, DMEM containing 5% FBS and 1% methylcellulose (Sigma) was overlaid to infected Vero cells for plaque formation. At 3 days post-infection, Vero monolayers were stained with 1% crystal violet for plaque counting.

### Mice

*Nampt^tm1Oleo^* mice (Jax stock *Nampt^tm1Oleo^*/ImaiJ; Strain #: 034242) were gifted from Prof. Shin-ichiro Imai at the Washington University School of Medicine.^41^ *Nampt^tm1Oleo^* mice were crossed with CreERT2 mice (Jax stock B6;129-*Gt(ROSA)26Sor^tm1(cre/ERT2)Tyi^*/J, Jax strain #: 008463) from Jackson Laboratories. Primers used in genotyping listed in Table S2. The PCR conditions are as follow: 20s at 95 °C, 30s at 55 °C, and 30s at 72 °C, for 30 cycles. To generate *Nampt^-/-^* mice, *Nampt^tm1Oleo^*/CreERT2 mice were injected intraperitoneally with tamoxifen in Kolliphor EL (75 mg/kg) for 5 days. To restore NAD^+^, 400 mg/kg of NR was supplemented daily via intraperitoneal injection.^42^ Age- (10-12 weeks) and gender-matched mice were used for all experiments, unless specified otherwise. All animal experiments were performed in accordance with the Guide for the Care and Use of Laboratory Animals of the NIH. The animal protocol was approved by the Institutional Animal Care and Use Committee (IACUC) of USC.

For HSV-1 infection, age- and gender-matched mice were challenged with HSV-1 (5 × 10^7^ PFU) in 100 μL of PBS via intravenous injection, at 3 days after tamoxifen treatment. Mouse survival was monitored daily for 20 days. To titrate viral replication and analyze immune response, infected mice were sacrificed at 3-day post-infection and tissues/organs were collected. One piece of the liver was fixed in 4% PFA for histology analysis, and the remaining was homogenized with Zirconia/Silica Beads by beats beater (Bio Spec Products Inc.) for immunoblotting, DNA/RNA extraction and virus titration. HSV-1 titer was determined by plaque assay, cytokine transcripts (after reverse transcription) and viral DNA were measured by real-time qRT-PCR. To evaluate the role of NR upon HSV-1 infection in mice, modest dose of HSV-1 (2 × 10^7^ PFU) was used in 100 μL of PBS for intravenous injection at 3 days after tamoxifen treatment. Mouse survival was monitored daily for 20 days, and tissues were collected for metabolite analysis, viral titration, immunoblotting and DNA/RNA extraction.

### NAMPT protein expression and purification

Human NAMPT (encoding residues 1-491) cDNAs, including wild-type, NAMPT-H247E and NAMPT-D219A, were cloned into the pGEX-4T-1 bacterial expression vector, with a TEV protease recognition site inserted after the GST tag. The NAMPT constructs were transformed into BL21 *E. coli* competent cells. After induction at 20 °C overnight, bacterial pellet was lysed, sonicated and centrifuged at 12,000 rpm (J-LITE JLA-16.250, Beckman) for 20 min at 4 °C. The cleared lysate was mixed with Glutathione Sepharose® 4B (Sigma, GE17-0756-01) and incubated at 4°C for 2 h with constant rotation. Glutathione beads were washed 3 times with 20 mM Tris pH 7.5, 0.1 % Triton X-100 and proteins were eluted with 10 mM reduced glutathione in 50 mM Tris pH 8.0. Purified protein was analyzed and quantified by SDS-PAGE and Coomassie staining and stored in 10% glycerol at -80°C for in vitro reactions.

### HSV-1 DNA extraction

To extract the viral DNA in supernatant, virions were pelleted via high-speed ultracentrifugation at 80,000 g for 1 h at 4°C. Viral DNA and cellular DNA were extracted as previously reported with minor modifications.^43^ In brief, the tissue, cell and virion samples were lysed in NK Lysis Buffer (50 mM Tris, 50 mM EDTA, 1% SDS, pH 8) with Proteinase K, and incubated at 55 °C overnight. RNase A was used to remove RNA. Samples were cooled on ice prior to the addition of pre-chilled 7.5 M ammonium acetate to precipitate proteins. After vortex, the samples were centrifuged at 4,000*g* for 10 minutes and the supernatant was carefully transferred into a new tube. To precipitate genomic DNA, 60% (v/v) isopropanol was added to the tube, followed by 20 times of inverting and centrifugation at 15, 000*g* for 10 minutes. DNA pellet was washed with freshly prepared 70% ethanol. After air drying for 10-30 minutes, 1 × TE buffer was added to fully dissolve the DNA pellet.

### RNA extraction and real-time qRT-PCR

Cellular total RNA was extracted using TRIzol reagent (Invitrogen) per manufacture’s instruction. Mouse tissue RNA was extracted using the RNeasy Mini Kit (Qiagen, 74106) following the manufacture’s instruction. The total of 1 μg RNA, treated with DNase I, was used as template to synthesize cDNA with M-MLV reverse transcriptase (Promega, M1701). Each quantitative real-time PCR (qRT-PCR) reaction was assembled with cDNA template, corresponding primer set (Table S2) and SYBR master mix (Applied Biosystems). Cycle conditions were as the following: 95 °C for 10 min and 95 °C for 15 s followed by 55 °C for 1 min (40 cycles). Relative mRNA expression for each target gene was calculated by the 2^-ΔΔCt^ method, normalized to the *Actb* for mouse or *ACTB* for human.

### Immunohistochemical, immunofluorescence analysis and Hematoxylin & Eosin (H&E) staining

To visualize the expression of NAMPT and viral proteins, extracted liver tissues were first fixed in 4% paraformaldehyde solution (Sigma) for 24 hours at 4 °C. After dehydration in 70%, 80%, 95%, 100% ethanol (v/v in water) and xylene, tissues were embedded in paraffin and sliced into 4-μm sections. For immunohistochemistry staining, the sections were deparaffinated, rehydrated, and oxidized. Processed sections were blocked with 10% goat serum for 1 hour, incubated with the primary antibodies at 4 °C overnight. After washing with PBST, sections were probed with horseradish peroxidase-linked secondary antibody (A120-101P, Bethyl Laboratories) for 40 min at 37 °C, reacted with diaminobenzidine (DAB, Leica) and counterstained with Mayer hematoxylin (Leica).

To perform H&E staining, deparaffinated sections were incubated with Hematoxylin for 30 s and rinsed with de-ionized water, followed by Eosin staining for 30 s, prior to dehydration and mounting. Each section was evaluated and captured under an inverted confocal microscope (Nikon Eclipse Ti2) equipped with 10-40× objective.

To determine the incorporation of NAMPT into HSV-1 virion particles, sgNAMPT HeLa cells stably reconstituted with NAMPT-HA were grown on poly-L-lysine-coated coverslips and infected with HSV-1. The cells were fixed in 4% paraformaldehyde and permeabilized with 0.1% Triton X-100. After blocking with 5% BSA in PBS, samples were incubated with primary antibodies at 4 °C overnight, followed by incubation with dye-conjugated secondary antibodies for one hour at room temperature. Imaging was performed using a Nikon eclipse Ti microscope with NIS-Elements AR imaging software.

### Sucrose gradient centrifugation

At 48 h post HSV-1 infection, the HSV-1-containing medium was collected, centrifuged at 4,000 rpm for 10 min, and filtered through 0.22 μM membrane. Virions were further pelleted by centrifugation at 80,000 g (Beckman TYPE45) for 60 min at 4°C and resuspended in TNE buffer (0.01 M Tris-HCl, 0.5 M NaCl, 1 mM EDTA, pH 7.5).^44,45^ Resuspended virions were loaded onto the top of the 15-55% (w/v) sucrose gradient and centrifuged at 80,000 g for 60 min at 4°C in a Beckman SW41 rotor. A total of 12 fractions, 1 mL sucrose solution for each, were collected and analyzed by SDS-PAGE and immunoblotting with antibodies against HSV-1 antigens. Virions were observed as milky bands identified by scattered light. HSV-1 virions were further pelleted down and resuspended in PBS or in vitro assay buffer (see in vitro phosphoribosylase activity assay) for further biochemical and microscopic analyses. Extraction of NAMPT exosomes (Flag-NAMPT) followed published protocols ^19,46^ and validated via immunoblotting with antibodies against CD81 (1:1000, Santa Cruz, sc-166029), Alix (1:1000, Santa Cruz, sc-53540), CD63 (1:1000, BD Pharmingen, car# 556019) and Calnexin (1:1000, Cell Signaling, C5C9).

### Co-sedimentation assay and proteinase K digestion

Proteinase K (final concentration of 50 μg/ml) or Triton X-100 (final concentration of 1%) was added to purified virion suspension and incubated for 5 min at 55°C. The proteinase activity was terminated by adding 5 mM phenylmethylsulfonyl fluoride for 10 minutes at room temperature. Alternatively, HSV-1 virions treated with NP40 buffer (1%) was incubated with recombinant NAMPT protein and centrifuged through sucrose gradient as previously described.^47^ Nuclear capsids were collected and analyzed by immunoblotting with antibodies against HSV-1 glycoprotein B, tegument UL37 and nucleocapsid UL19 protein.

### Immunogold staining and transmission electron microscopy

To prepare virions for transmission electron microscopy (TEM), HSV-1 virions were purified from sgNAMPT HeLa cells reconstituted with HA-NAMPT. Purified viral particles were pretreated with 2% Triton X-100 and 2% glutaraldehyde for 30 min. Viral particles were adsorbed on a formvar-carbon coated nickel grids for 30 min. To prepare cell sections for transmission electron microscopy, cells were fixed with 4% paraformaldehyde and 0.05% glutaraldehyde in PIPES buffer for one hour on ice. Cells were dehydrated, sectioned and placed on grids.

For immunogold labelling, the samples were incubated in blocking buffer (PBS, 1% BSA, 1% Tween-20) for one hour, followed by the incubation with primary antibodies (anti-HA/NAMPT, rabbit polyclonal HA antibody, BioLegend Cat #: 901501, RRID: AB_2565006, dilution: 1:100; anti-UL19, mouse monoclonal [3B6] to HSV1 + HSV2 UL19 major capsid protein, Abcam Cat #: ab6508, RRID: AB_305530, dilution 1:100; anti-TGN46, Sigma Cat #: ABT95) at 4 °C overnight and gold-conjugated secondary antibodies at room temperature for one hour. After being rinsed with fresh PBS containing 1% Tween-20, grids were stained with 1% uranyl acetate (10 min at room temperature), followed by 0.25% lead citrate (2 min at room temperature), and air dried. All electron microscopy analyses were carried out on a JEOL 2100 transmission electron microscope operated at 50 keV.

### Identification of phosphoribosylated peptides by mass spectrometry

Phosphoribosylated peptides were analyzed on the EASY-nLC 1200 coupled to an Orbitrap Eclipse Tribrid mass spectrometer (ThermoFisher Scientific, San Jose, CA). Peptides were separated on an Aurora UHPLC Column (25 cm × 75 μm, 1.6 μm C18, AUR2-25075C18A, Ion Optics) with a flow rate of 0.35 μL/min for a total duration of 135 min and ionized at 1.6 kV in the positive ion mode. The gradient was composed of 6% solvent B (7.5 min), 6-25% B (82.5 min), 25-40% B (30 min), and 40–98% B (15 min); solvent A: 0.1% formic acid in water; solvent B: 80% ACN and 0.1% formic acid. MS1 scans were acquired at the resolution of 120,000 from 350 to 2,000 m/z, AGC target 1e6, and maximum injection time 50 ms. MS2 scans were obtained in the ion trap using fast scan rate on precursors with 2-7 charge states and quadrupole isolation mode (isolation window: 0.7 m/z) with higher-energy collisional dissociation (HCD, 30%) activation type. Dynamic exclusion was set to 30 s. The temperature of ion transfer tube was 300°C and the S-lens RF level was set to 30. MS2 fragmentation spectra were searched with Proteome Discoverer SEQUEST (version 2.5, Thermo Scientific) against in silico tryptic digested Uniprot Human herpesvirus 1 (HHV-1) database. The maximum missed cleavages were set to 2. Dynamic modifications were set to oxidation on methionine (M, +15.995 Da), phosphoribosylation (D, E, R and K, +212.009 Da), deamidation (N and Q, +0.984 Da), protein N-terminal acetylation (+42.011 Da) and Met-loss (-131.040 Da). Carbamidomethylation on cysteine residues (C, +57.021 Da) was set as a fixed modification. The maximum parental mass error was set to 10 ppm, and the MS2 mass tolerance was set to 0.6 Da. The false discovery threshold was set strictly to 0.01 using the Percolator Node validated by q-value. The relative abundance of parental peptides was calculated by integration of the area under the curve of the MS1 peaks using the Minora LFQ node.

### Metabolite analysis by liquid chromatography-mass spectrometry

Mock- and HSV-1-infected cells (approximately 2×10^6^) were first washed with PBS and ice-cold 150 mM ammonium acetate (NH4AcO, pH 7.3). One mL of 80% MeOH, pre-chilled at -80°C, was added to cells to extract metabolites. After incubation at -80°C for 20 mins, cells were scraped off and pelleted at 4°C for 5 min at 21,000 g. The supernatant was transferred into new microfuge tubes, vacuum-dried at room temperature; and the pellet was re-suspended in water for LC-MS analysis. Samples from in vitro assays, were centrifuged to separate protein and the supernatant. The supernatant was mixed with cold absolute MeOH to extract metabolites and vacuum dried as described above. Resuspended samples were randomized and analyzed on a Q-Exactive Plus hybrid quadrupole-Orbitrap MS coupled with Vanquish UHPLC system (ThermoFisher Scientific) in polarity switching mode (+3.00 kV/-2.25 kV) with an m/z window ranging from 65 to 975. Mobile phase A was 5 mM NH4AcO, pH 9.9, and mobile phase B was acetonitrile. Metabolites were separated on a Luna 3 μm NH2 100 NH2 100A° (150 x 2.0 mm) column (Phenomenex), with the flow rate of 0.3 mL/min, gradient from 15% A to 95% A in 18 min, followed by an isocratic step for 9 min and re-equilibration for 7 min. Each metabolite was identified based on retention time and standard compound (Sigma) and quantified by peak area integration using the TraceFinder 4.1 (ThermoFisher Scientific).

### Two-dimensional gel electrophoresis

Protein samples, including whole cell lysates, purified virions and in vitro phosphoribosylase assay samples, were resolved in 150 μL rehydration buffer (8 M urea, 2% CHAPS, 0.5% IPG buffer, and 0.002% bromophenol blue), followed by sonication for 20 s and centrifugation at 20,000 g for 2 min. The supernatant was incubated with 6-11NL or 3-11NL IEF dry strips with a running conditions: 20 V, 10 h; hold at 100 V, 1 h; 500 V, 1 h and 1,000 V, 1 h; gradient mode at 5,000 V for 4 h and hold at 5,000 V for 4 h. Prior to SDS-PAGE, strips were incubated with SDS-equilibration buffer (50mM Tris-HCl [pH 8.8], 6M urea, 30% glycerol, 2% SDS, 0.001% bromophenol blue) containing 10 mg/ml DTT for 15 min and that containing 25 mg/ml 2-iodoacetamide for 15 min. Strips were rinsed with SDS-PAGE buffer, resolved by SDS-PAGE, and analyzed by immunoblotting with indicated primary antibodies. Primary antibodies used to probe GST or epitopes (V5, Flag, or HA) for HSV-1 proteins include mouse monoclonal GST antibody (Santa Cruz Biotechnology Cat#: sc-138) diluted at 1:1,000, monoclonal Flag M2 monoclonal antibody (Sigma Cat #: F1804) diluted at 1:2,000; rabbit polyclonal V5 antibody (Bethyl Cat#: A190-120A, RRID: AB_67586) diluted at 1:1,000.

### NAMPT phosphoribosyltransferase assay in the salvage NAD^+^ synthesis

NAMPT activity was assessed using CycLex® NAMPT Colorimetric Assay Kit Ver.2 (MBL, CY-1251V2) per manufacturer’s instructions. Briefly, purified wild-type and mutant NAMPT (tagged with either Streptavidin-Binding Peptide (SBP) or GST) mixed with NAM, PRPP, ATP, diaphorase, ADH, NMNAT1, WST-1 and ethanol in the Tris-HCl and MgCl2 buffer. The mixture was incubated at 30°C and the absorbance at 450 nm was determined to represent the enzymatic activity.

### In vitro NAMPT protein phosphoribosylase assay

The in vitro protein phosphoribosylase assay was performed as previously described with modifications.^48,49^ In brief, 5 μg of purified GST-VP22 or virions (∼10 μg total viral proteins) were incubated with 1 μg of purified NAMPT or its mutants at 37°C for 30 min in NAMPT reaction buffer (50 mM Tris-HCl [pH 7.5], 2 mM DTT, 10 mM MgCl2, and 2 mM ATP). The reaction was stopped by adding rehydration buffer. VP22 was eluted and analyzed by two-dimensional gel electrophoresis and immunoblotting with corresponding antibodies.

To quantitatively determine phosphoribose released by NAMPT, up to 20 μg of purified HSV-1 virions, and 2-fold dilutions were incubated with 1 μg of purified GST-NAMPT or its mutants at 37°C for 30 min in NAMPT reaction buffer. Before in vitro reaction, purified HSV-1 virions were extensively washed with in vitro reaction buffer to reduce background of released phosphoribose. Reaction samples were centrifuged at 80,000 g to pellet HSV-1 virions. The supernatant (80 μL) was mixed with 640 μL methanol (LC/MS/MS grade) to extract metabolites that were subjected to LC-MS analysis.

### Co-immunoprecipitation (Co-IP)

HEK293T cells were infected with HSV-1 or transfected with indicated plasmids for 36-48 hours. Whole cell lysates were prepared with NP40 buffer (50 mM Tris-HCl, pH 7.4, 150 mM NaCl, 1% NP-40, 5 mM EDTA) supplemented with 20 mM β-glycerophosphate and 1 mM sodium orthovanadate. Samples were sonicated, centrifuged and pre-cleared with Sepharose® 4B agarose beads (Sigma) for 1 hour at 4 °C. Pre-cleared samples were incubated with ANTI-FLAG® M2 Agarose Affinity Gel or Anti-V5 Agarose Affinity Gel (Bethyl Group) for 4 hours at 4 °C. The agarose beads were washed extensively, and samples were eluted by boiling at 95 °C for 10 min. Precipitated proteins were analyzed by SDS-PAGE and immunoblotting.

### SDS-PAGE and immunoblotting

Tissues and cells were solubilized in lysis buffer (50 mM Tris-HCl, pH 7.4, 1 mM EDTA, 150 mM NaCl, 0.25% Na-deoxycholate, 1% NP-40, 0.10% SDS, 1% TritonX-100 on ice for 30 min and lysates were centrifuged at 12000 rpm at 4 °C for 20 min. Soluble fractions were denatured in loading buffer at 95 °C and separated by SDS-PAGE and subsequently transferred to nitrocellulose membranes. Membranes were blocked with 5% skim milk and probed with primary antibodies overnight at 4 °C, later incubated with secondary antibodies (IRDye800-conjugated 1: 10,000 dilutions, LI-COR Biosciences) for 1 hour before exposed on the LI-COR Odyssey infrared imager. Images were processed using Image Studio Version 4.0 (LI-COR Biosciences). Primary antibodies (diluted 1:1,000) were used in this study are listed in the KEY

## RESOURCES TABLE

### Engineering recombinant HSV-1

HSV-1 mutant viruses used in this study were constructed with an infectious clone of pBAC-HSV-1.^50^ Mutagenesis was created based on a markerless two-step Red recombination protocol.^51^ In brief, linear PCR products, with a Kan cassette contained an *I-Sce*I homing endonuclease site and mutation site(s), were amplified using the pEPkan-S vector and primers with 60 bp homology to the region of interest. Purified PCR products were electroporated into GS1783 bacteria containing pBAC-HSV-1. The first lambda Red recombination incorporated the PCR product into the region of interest via homologous recombination. The second lambda Red recombination step was performed to remove the *Kan^R^*-*I-Sce*I gene cassette, by utilizing an *I-Sce*I homing endonuclease and the repeat sequence within the homologous arms of the inserted PCR products. Engineered pBAC-HSV-1 plasmids were extracted using isopropanol precipitation method. The mutations was confirmed by PCR and sequencing. The primer sequences are listed in Table S2.

To generate mutant HSV-1, approximately 1 × 10^6^ Vero cells were transfected with 2 μg of extracted pBAC-HSV-1 DNA using Lipofectamine 3000 (Invitrogen). In general, viral plaques were observed at 48 to 72 h post-transfection. For some recombinant HSV-1 that lytic replication is reduced, cells were splitted to allow cytopathic effect to appear. Recombinant HSV-1 was subjected to two rounds of plaque purification, followed by three to four times of amplification to produce sufficient viruses for biochemical and virological experiments.

### HSV-1 entry assay

NAMPT-treated or mutant HSV-1 virions were diluted in FBS-free DMEM and used to infect HeLa cells for up to 2 hours. Infected cells were disassociated by trypsin digestion and washed extensively with PBS. Total genomic DNA was extracted, and the HSV-1 genome was quantified by real-time PCR using primers specific to HSV-1 genes, i.e., UL48 and UL37. To generate HSV-1 virions containing mutant viral proteins (such as VP22, gB, gD and gH), plasmids containing wild-type or mutant VP22, gD, gB and gH were first transfected into HEK293T cells. At 24 h post-transfection, cells were infected with wild-type HSV-1 at MOI=0.1. HSV-1 progeny virions in the medium were collected and normalized by quantitative PCR for viral genome. Equal amount of HSV-1 virions were used for entry assay as described above.

## QUANTIFICATION AND STATISTICAL ANALYSIS

### Statistical analysis

Data represent the mean of three independent experiments, and error bars denote standard deviation unless specified otherwise. GraphPad Prism 9 was used to analyze the mouse Kaplan-Meier survival curve, the log-rank test applied to the mouse survival data and Michaelis-Menten modeling to calculate Vmax, Kcat and Km. Statistics was also analyzed with GraphPad Prism 9 for unpaired two-tailed Student’s *t*-test and repeated-measures of ANOVA. A p-value less than 0.05 is considered statistically significant. *, *P<0.05; **, P<0.01; ***, P<0.001*.

## References

1 Girdhar, K., Powis, A., Raisingani, A., Chrudinova, M., Huang, R., Tran, T., Sevgi, K., Dogus Dogru, Y. & Altindis, E. (2021). Viruses and Metabolism: The Effects of Viral Infections and Viral Insulins on Host Metabolism. Annu Rev Virol 8, 373–391, 10.1146/annurev-virology-091919-102416.

2 Schneider, W. M., Chevillotte, M. D. & Rice, C. M. (2014). Interferon-stimulated genes: a complex web of host defenses. Annu Rev Immunol 32, 513–545, 10.1146/annurev-immunol-032713-120231.

3 Sadler, A. J. & Williams, B. R. (2008). Interferon-inducible antiviral effectors. Nat Rev Immunol 8, 559–568, 10.1038/nri2314.

4 Lahouassa, H., Daddacha, W., Hofmann, H., Ayinde, D., Logue, E. C., Dragin, L., Bloch, N., Maudet, C., Bertrand, M., Gramberg, T. et al. (2012). SAMHD1 restricts the replication of human immunodeficiency virus type 1 by depleting the intracellular pool of deoxynucleoside triphosphates. Nat Immunol 13, 223–228, 10.1038/ni.2236.

5 Gizzi, A. S., Grove, T. L., Arnold, J. J., Jose, J., Jangra, R. K., Garforth, S. J., Du, Q., Cahill, S. M., Dulyaninova, N. G., Love, J. D. et al. (2018). A naturally occurring antiviral ribonucleotide encoded by the human genome. Nature 558, 610–614, 10.1038/s41586-018-0238-4.

6 Liu, S. Y., Aliyari, R., Chikere, K., Li, G., Marsden, M. D., Smith, J. K., Pernet, O., Guo, H., Nusbaum, R., Zack, J. A. et al. (2013). Interferon-inducible cholesterol-25-hydroxylase broadly inhibits viral entry by production of 25-hydroxycholesterol. Immunity 38, 92–105, 10.1016/j.immuni.2012.11.005.

7 Verdin, E. (2015). NAD(+) in aging, metabolism, and neurodegeneration. Science 350, 1208–1213, 10.1126/science.aac4854.

8 Yoshino, J., Baur, J. A. & Imai, S. I. (2018). NAD(+) Intermediates: The Biology and Therapeutic Potential of NMN and NR. Cell Metab 27, 513–528, 10.1016/j.cmet.2017.11.002.

9 Lautrup, S., Sinclair, D. A., Mattson, M. P. & Fang, E. F. (2019). NAD(+) in Brain Aging and Neurodegenerative Disorders. Cell Metab 30, 630–655, 10.1016/j.cmet.2019.09.001.

10 Huang, D. & Kraus, W. L. (2022). The expanding universe of PARP1-mediated molecular and therapeutic mechanisms. Mol Cell 82, 2315–2334, 10.1016/j.molcel.2022.02.021.

11 Groslambert, J., Prokhorova, E. & Ahel, I. (2021). ADP-ribosylation of DNA and RNA. DNA Repair (Amst) 105, 103144, 10.1016/j.dnarep.2021.103144.

12 Challa, S., Khulpateea, B. R., Nandu, T., Camacho, C. V., Ryu, K. W., Chen, H., Peng, Y., Lea, J. S. & Kraus, W. L. (2021). Ribosome ADP-ribosylation inhibits translation and maintains proteostasis in cancers. Cell 184, 4531–4546 e4526, 10.1016/j.cell.2021.07.005.

13 Palazzo, L., Daniels, C. M., Nettleship, J. E., Rahman, N., McPherson, R. L., Ong, S. E., Kato, K., Nureki, O., Leung, A. K. & Ahel, I. (2016). ENPP1 processes protein ADP-ribosylation in vitro. FEBS J 283, 3371–3388, 10.1111/febs.13811.

14 Thirawatananond, P., McPherson, R. L., Malhi, J., Nathan, S., Lambrecht, M. J., Brichacek, M., Hergenrother, P. J., Leung, A. K. L. & Gabelli, S. B. (2019). Structural analyses of NudT16-ADP-ribose complexes direct rational design of mutants with improved processing of poly(ADP-ribosyl)ated proteins. Sci Rep 9, 5940, 10.1038/s41598-019-39491-w.

15 O’Sullivan, J., Tedim Ferreira, M., Gagne, J. P., Sharma, A. K., Hendzel, M. J., Masson, J. Y. & Poirier, G. G. (2019). Emerging roles of eraser enzymes in the dynamic control of protein ADP-ribosylation. Nat Commun 10, 1182, 10.1038/s41467-019-08859-x.

16 Katsyuba, E., Romani, M., Hofer, D. & Auwerx, J. (2020). NAD(+) homeostasis in health and disease. Nat Metab 2, 9–31, 10.1038/s42255-019-0161-5.

17. 17 Garten, A., Schuster, S., Penke, M., Gorski, T., de Giorgis, T. & Kiess, W. (2015). Physiological and pathophysiological roles of NAMPT and NAD metabolism. Nat Rev Endocrinol 11, 535–546, 10.1038/nrendo.2015.117.

18 Schoggins, J. W., Wilson, S. J., Panis, M., Murphy, M. Y., Jones, C. T., Bieniasz, P. & Rice, C. M. (2011). A diverse range of gene products are effectors of the type I interferon antiviral response. Nature 472, 481–485, 10.1038/nature09907.

19 Yoshida, M., Satoh, A., Lin, J. B., Mills, K. F., Sasaki, Y., Rensing, N., Wong, M., Apte, R. S. & Imai, S. I. (2019). Extracellular Vesicle-Contained eNAMPT Delays Aging and Extends Lifespan in Mice. Cell Metab 30, 329–342 e325, 10.1016/j.cmet.2019.05.015.

20 Smith, G. A. (2017). Assembly and Egress of an Alphaherpesvirus Clockwork. Adv Anat Embryol Cell Biol 223, 171–193, 10.1007/978-3-319-53168-7_8.

21. 21 Laine, R. F., Albecka, A., van de Linde, S., Rees, E. J., Crump, C. M. & Kaminski, C. F. (2015). Structural analysis of herpes simplex virus by optical super-resolution imaging. Nat Commun 6, 5980, 10.1038/ncomms6980.

22 Stremlau, M., Owens, C. M., Perron, M. J., Kiessling, M., Autissier, P. & Sodroski, J. (2004). The cytoplasmic body component TRIM5alpha restricts HIV-1 infection in Old World monkeys. Nature 427, 848–853, 10.1038/nature02343.

23 Dasgupta, S., Rajapakshe, K., Zhu, B., Nikolai, B. C., Yi, P., Putluri, N., Choi, J. M., Jung, S. Y., Coarfa, C., Westbrook, T. F. et al. (2018). Metabolic enzyme PFKFB4 activates transcriptional coactivator SRC-3 to drive breast cancer. Nature 556, 249–254, 10.1038/s41586-018-0018-1.

24 Zhao, J., Tian, M., Zhang, S., Delfarah, A., Gao, R., Rao, Y., Savas, A. C., Lu, A., Bubb, L., Lei, X. et al. (2020). Deamidation Shunts RelA from Mediating Inflammation to Aerobic Glycolysis. Cell Metab 31, 937–955 e937, 10.1016/j.cmet.2020.04.006.

25 Wang, T., Zhang, X., Bheda, P., Revollo, J. R., Imai, S. & Wolberger, C. (2006). Structure of Nampt/PBEF/visfatin, a mammalian NAD+ biosynthetic enzyme. Nat Struct Mol Biol 13, 661–662, 10.1038/nsmb1114.

26 Daniels, C. M., Ong, S. E. & Leung, A. K. L. (2017). ADP-Ribosylated Peptide Enrichment and Site Identification: The Phosphodiesterase-Based Method. Methods Mol Biol 1608, 79–93, 10.1007/978-1-4939-6993-7_7.

27 Matic, I., Ahel, I. & Hay, R. T. (2012). Reanalysis of phosphoproteomics data uncovers ADP-ribosylation sites. Nat Methods 9, 771–772, 10.1038/nmeth.2106.

28 Zhang, Y., Wang, J., Ding, M. & Yu, Y. (2013). Site-specific characterization of the Asp- and Glu-ADP-ribosylated proteome. Nat Methods 10, 981–984, 10.1038/nmeth.2603.

29 Revollo, J. R., Grimm, A. A. & Imai, S. (2004). The NAD biosynthesis pathway mediated by nicotinamide phosphoribosyltransferase regulates Sir2 activity in mammalian cells. J Biol Chem 279, 50754–50763, 10.1074/jbc.M408388200.

30 Connolly, S. A., Jardetzky, T. S. & Longnecker, R. (2021). The structural basis of herpesvirus entry. Nat Rev Microbiol 19, 110–121, 10.1038/s41579-020-00448-w.

31 Chesnokova, L. S., Jiang, R. & Hutt-Fletcher, L. M. (2015). Viral Entry. Curr Top Microbiol Immunol 391, 221–235, 10.1007/978-3-319-22834-1_7.

32. 32 Van den Bergh, R., Florence, E., Vlieghe, E., Boonefaes, T., Grooten, J., Houthuys, E., Tran, H. T., Gali, Y., De Baetselier, P., Vanham, G., et al. (2010). Transcriptome analysis of monocyte-HIV interactions. Retrovirology 7, 53, 10.1186/1742-4690-7-53.

33 Li, J., Zhao, J., Xu, S., Zhang, S., Zhang, J., Xiao, J., Gao, R., Tian, M., Zeng, Y., Lee, K. et al. (2019). Antiviral activity of a purine synthesis enzyme reveals a key role of deamidation in regulating protein nuclear import. Sci Adv 5, eaaw7373, 10.1126/sciadv.aaw7373.

34 He, S., Zhao, J., Song, S., He, X., Minassian, A., Zhou, Y., Zhang, J., Brulois, K., Wang, Y., Cabo, J. et al. (2015). Viral Pseudo-Enzymes Activate RIG-I via Deamidation to Evade Cytokine Production. Mol Cell, 10.1016/j.molcel.2015.01.036.

35 Pan, C., Li, B. & Simon, M. C. (2021). Moonlighting functions of metabolic enzymes and metabolites in cancer. Mol Cell 81, 3760–3774, 10.1016/j.molcel.2021.08.031.

36 Weixler, L., Scharinger, K., Momoh, J., Luscher, B., Feijs, K. L. H. & Zaja, R. (2021). ADP-ribosylation of RNA and DNA: from in vitro characterization to in vivo function. Nucleic Acids Res 49, 3634–3650, 10.1093/nar/gkab136.

37 Munnur, D., Bartlett, E., Mikolcevic, P., Kirby, I. T., Rack, J. G. M., Mikoc, A., Cohen, M. S. & Ahel, I. (2019). Reversible ADP-ribosylation of RNA. Nucleic Acids Res 47, 5658–5669, 10.1093/nar/gkz305.

38 Wang, X., Zhang, Q., Bao, R., Zhang, N., Wang, Y., Polo-Parada, L., Tarim, A., Alemifar, A., Han, X., Wilkins, H. M. et al. (2017). Deletion of Nampt in Projection Neurons of Adult Mice Leads to Motor Dysfunction, Neurodegeneration, and Death. Cell Rep 20, 2184–2200, 10.1016/j.celrep.2017.08.022.

39 Zhang, L. Q., Van Haandel, L., Xiong, M., Huang, P., Heruth, D. P., Bi, C., Gaedigk, R., Jiang, X., Li, D. Y., Wyckoff, G., et al. (2017). Metabolic and molecular insights into an essential role of nicotinamide phosphoribosyltransferase. Cell Death Dis 8, e2705, 10.1038/cddis.2017.132.

40 Bhogaraju, S., Kalayil, S., Liu, Y., Bonn, F., Colby, T., Matic, I. & Dikic, I. (2016). Phosphoribosylation of Ubiquitin Promotes Serine Ubiquitination and Impairs Conventional Ubiquitination. Cell 167, 1636–1649 e1613, 10.1016/j.cell.2016.11.019.

41 Yoon, M. J., Yoshida, M., Johnson, S., Takikawa, A., Usui, I., Tobe, K., Nakagawa, T., Yoshino, J. & Imai, S. (2015). SIRT1-Mediated eNAMPT Secretion from Adipose Tissue Regulates Hypothalamic NAD+ and Function in Mice. Cell Metab 21, 706–717, 10.1016/j.cmet.2015.04.002.

42 Trammell, S. A., Schmidt, M. S., Weidemann, B. J., Redpath, P., Jaksch, F., Dellinger, R. W., Li, Z., Abel, E. D., Migaud, M. E. & Brenner, C. (2016). Nicotinamide riboside is uniquely and orally bioavailable in mice and humans. Nat Commun 7, 12948, 10.1038/ncomms12948.

43 Chen, S., Sanjana, N. E., Zheng, K., Shalem, O., Lee, K., Shi, X., Scott, D. A., Song, J., Pan, J. Q., Weissleder, R. et al. (2015). Genome-wide CRISPR screen in a mouse model of tumor growth and metastasis. Cell 160, 1246–1260, 10.1016/j.cell.2015.02.038.

44 Dai, X. & Zhou, Z. H. (2014). Purification of Herpesvirus Virions and Capsids. Bio Protoc 4, 10.21769/bioprotoc.1193.

45 Cardone, G., Newcomb, W. W., Cheng, N., Wingfield, P. T., Trus, B. L., Brown, J. C. & Steven, A. C. (2012). The UL36 tegument protein of herpes simplex virus 1 has a composite binding site at the capsid vertices. J Virol 86, 4058–4064, 10.1128/JVI.00012-12.

46 Cheng, Q., Shi, X. & Zhang, Y. (2020). Reprogramming Exosomes for Immunotherapy. Methods Mol Biol 2097, 197–209, 10.1007/978-1-0716-0203-4_12.

47 Langelier, C. R., Sandrin, V., Eckert, D. M., Christensen, D. E., Chandrasekaran, V., Alam, S. L., Aiken, C., Olsen, J. C., Kar, A. K., Sodroski, J. G. et al. (2008). Biochemical characterization of a recombinant TRIM5alpha protein that restricts human immunodeficiency virus type 1 replication. J Virol 82, 11682–11694, 10.1128/JVI.01562-08.

48 Zhang, R. Y., Qin, Y., Lv, X. Q., Wang, P., Xu, T. Y., Zhang, L. & Miao, C. Y. (2011). A fluorometric assay for high-throughput screening targeting nicotinamide phosphoribosyltransferase. Anal Biochem 412, 18–25, 10.1016/j.ab.2010.12.035.

49 Gardell, S. J., Hopf, M., Khan, A., Dispagna, M., Hampton Sessions, E., Falter, R., Kapoor, N., Brooks, J., Culver, J., Petucci, C. et al. (2019). Boosting NAD(+) with a small molecule that activates NAMPT. Nat Commun 10, 3241, 10.1038/s41467-019-11078-z.

50 Cunningham, C. & Davison, A. J. (1993). A cosmid-based system for constructing mutants of herpes simplex virus type 1. Virology 197, 116–124, 10.1006/viro.1993.1572.

51 Brulois, K. F., Chang, H., Lee, A. S., Ensser, A., Wong, L. Y., Toth, Z., Lee, S. H., Lee, H. R., Myoung, J., Ganem, D. et al. (2012). Construction and manipulation of a new Kaposi’s sarcoma-associated herpesvirus bacterial artificial chromosome clone. J Virol 86, 9708–9720, 10.1128/JVI.01019-12.

