## Supplemental figures for "The Interferon-inducible NAMPT acts as a protein phosphoribosylase to restrict viral infection"

**Affiliations:**
[1]Section of Infection and Immunity, Herman Ostrow School of Dentistry, Norris Comprehensive Cancer Center, University of Southern California, Los Angeles, CA 90089, USA;
[2]State Key Laboratory of Biotherapy and Cancer Center, West China Hospital and West China, School of Basic Medical Sciences & Forensic Medicine, Sichuan University, and Collaborative Innovation Center for Biotherapy, Chengdu, Sichuan 610041, P.R. China;
[3]Proteome Exploration Laboratory, Beckman Institute, California Institute of Technology, Pasadena, CA 91125, USA;
[4]Department of Diabetes and Cancer Metabolism, Beckman Research Institute, City of Hope National Medical Center, Duarte, CA 91010, USA.

† These authors contributed equally.
[#]Current address: Department of Diabetes and Cancer Metabolism, Beckman Research Institute, City of Hope National Medical Center, Duarte, CA 91010, USA.

### Figure S1

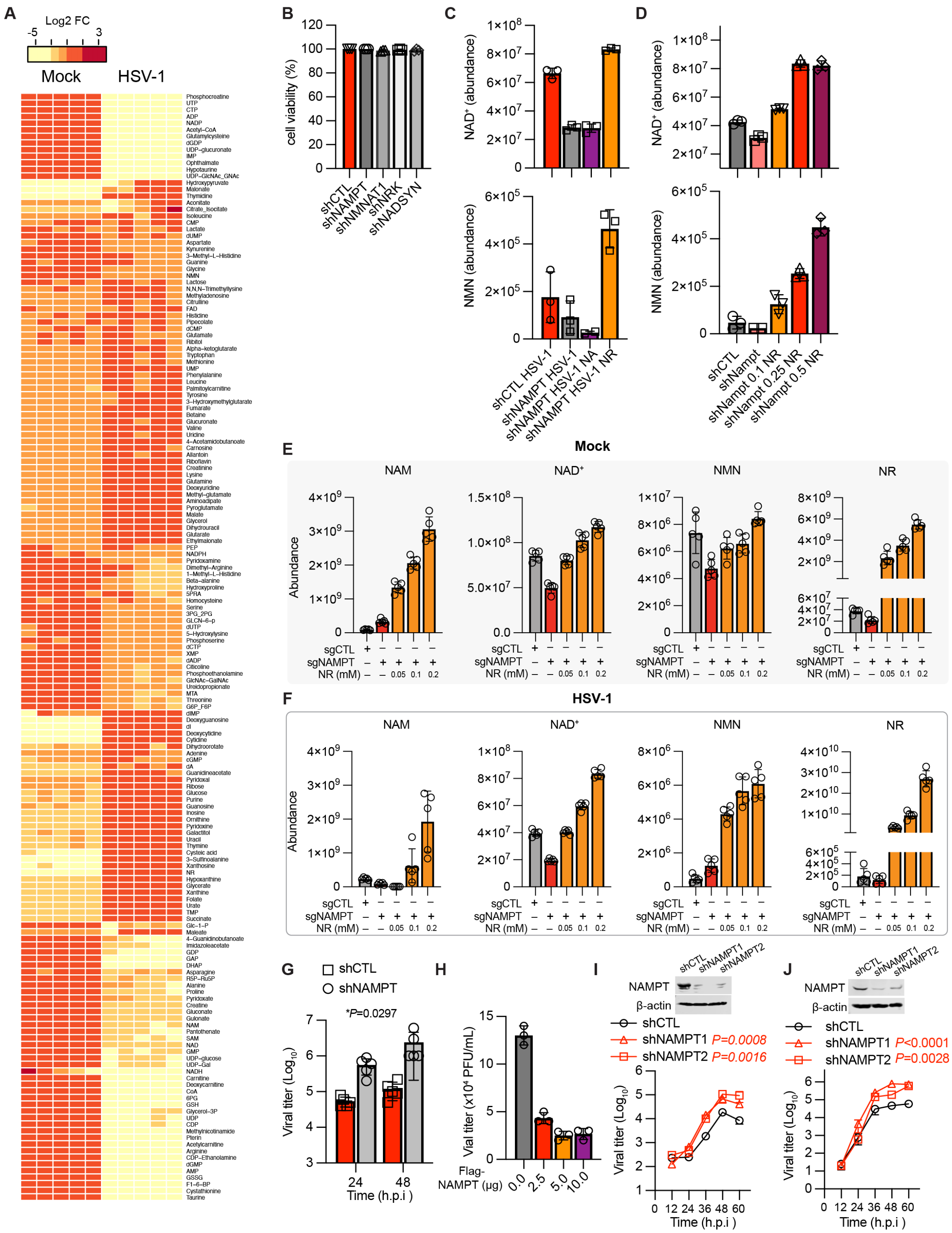

A
Log2 FC
Mock
HSV-1
B
C
D
E
Mock
NAM
NAD⁺
NMN
NR
Abundance
F
HSV-1
G
shCTL
shNAMPT
*P=0.0297
Viral titer (Log10)
Time (h.p.i )
H
Vial titer (x10⁴ PFU/mL)
Flag-NAMPT (μg)
I
NAMPT
β-actin
shCTL
shNAMPT1 P=0.0008
shNAMPT2 P=0.0016
J
shNAMPT1 P<0.0001
shNAMPT2 P=0.0028

**Figure S1. NAMPT restricts HSV-1 infection in cells independent of $NAD^+$**

(A) Heatmap of metabolites profiled in mock- and HSV-1-infected HepG2 cells at 12 hours post-infection (h.p.i) (MOI=5).

(B) Cell viability of 293T cells depleted with NAMPT, NMNAT1, NRK and NADSYN.

(C) Quantification of $NAD^+$ and NMN in shNAMPT HeLa cells infected with HSV-1, supplemented with nicotinic acid (NA, 0.1 mM) or nicotinamide riboside (NR, 0.1 mM).

(D) NR (0.1 mM) restores $NAD^+$ and NMN in shNAMPT HeLa cells infected with HSV-1.

(E and F) Exogenous NR increases the abundance of the $NAD^+$-related metabolites in mock- (E) and HSV-1-infected (F) sgNAMPT HeLa cells in a dose-dependent manner.

(G) HSV-1 titer in the medium of shCTL and shNAMPT HepG2 cells (supplemented with NR at 0.1 mM) at MOI=0.1 determined by plaque assay.

(H) HSV-1 titer in the medium of 293T cells that transiently express Flag-NAMPT in an increasing dose as indicated by plasmid amount at 24 h.p.i (MOI=0.1).

(I and J) Growth curve of HSV-1 in the medium of shCTL and shNAMPT mouse embryonic fibroblasts (MEFs) (I) and human foreskin fibroblasts (HFF) (J) at MOI=0.1, with NAMPT knockdown validated by immunoblotting with indicated antibodies using whole cell lysates.

Statistical significance was calculated using two-way ANOVA.

Related to Figure 1.

Figure S2

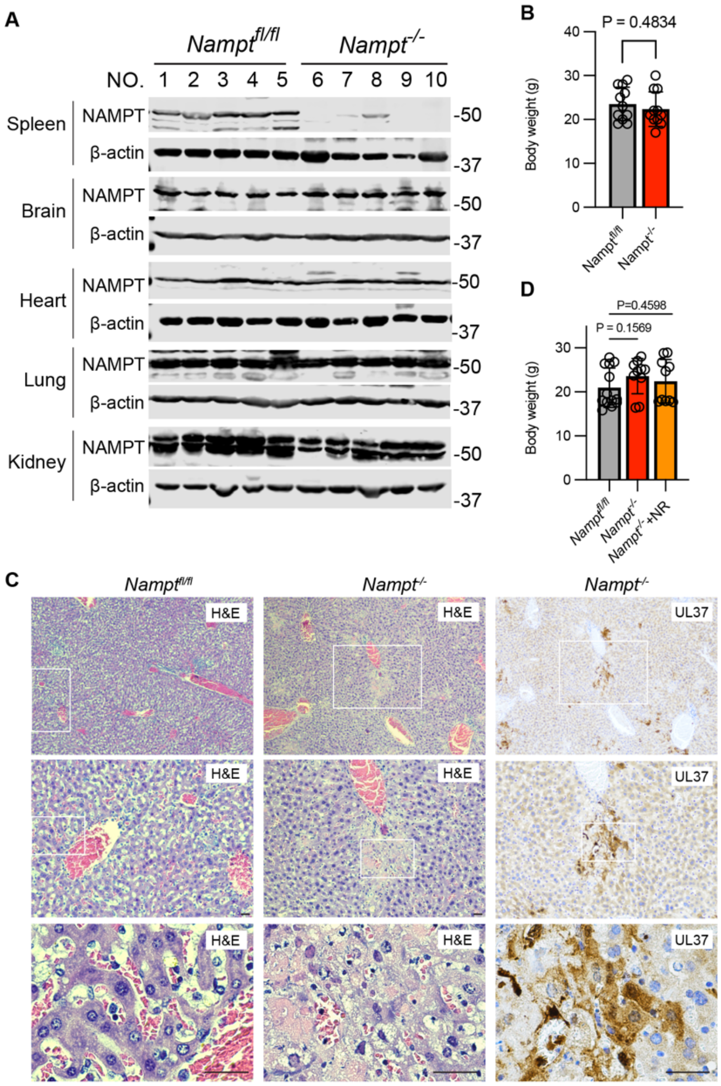


**Figure S2. NAMPT restricts HSV-1 infection in mice**

(A) Lysates of indicated tissues of wildtype and NAMPT-knockout (KO) mice were analyzed by immunoblotting with indicated antibodies.

(B) Body weight of wildtype and NAMPT-KO were determined right before HSV-1 infection.

(C) Hematolysin & Eosin (H&E) staining and UL37 immunohistochemistry staining of the liver of HSV-1-infected *Nampt*$^{fl/fl}$ and *Nampt*$^{-/-}$ (KO) mice, with boxed region shown below.

(D) Body weight of wildtype, NAMPT-KO and NAMPT-KO mice with NR injection were determined immediately before HSV-1 infection.

Related to Figure 2.

Figure S3

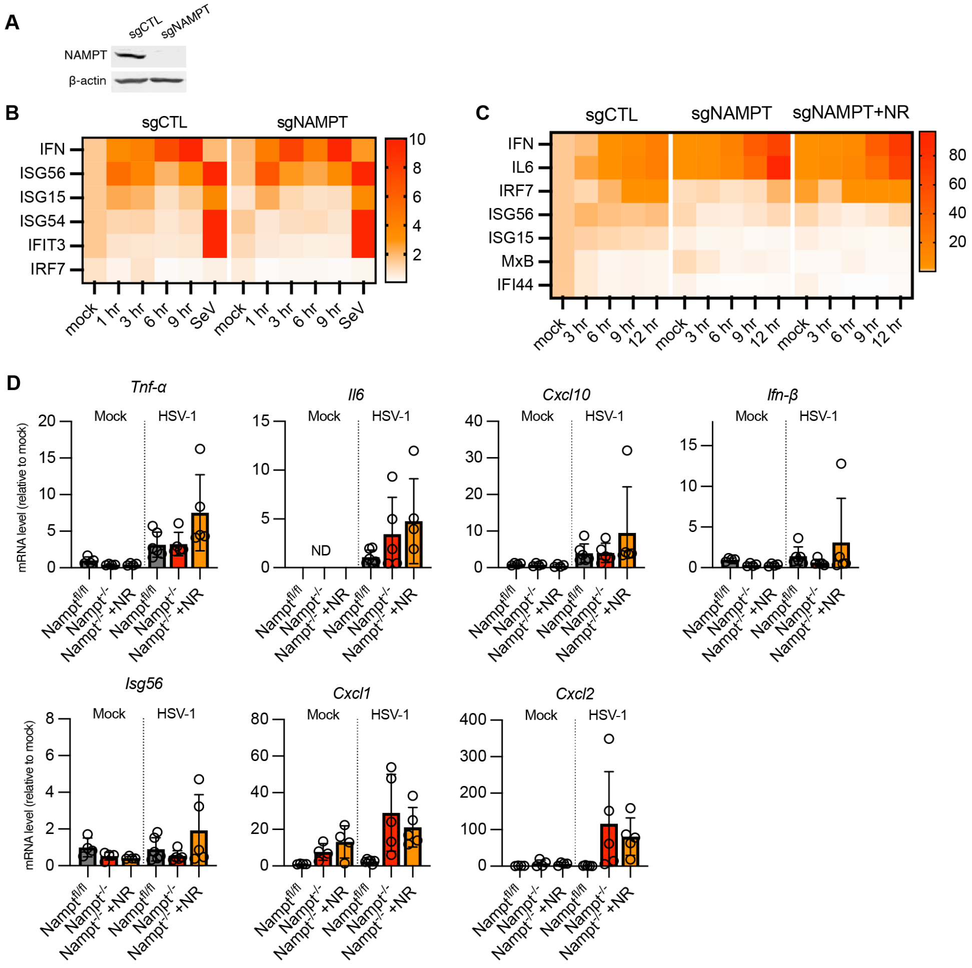


**Figure S3. Loss of NAMPT does not compromise innate immune response against HSV-1.**

(A) NAMPT knockout in HepG2 cells was validated by immunoblotting using whole cell lysates with indicated antibodies.

(B and C) Heatmap of cytokine gene expression in sgCTL and sgNAMPT HepG2, with Sendai virus (SeV, 30 HA unit/ml) (B), and sgCTL, sgNAMPT and sgNAMPT HeLa cells with NR (0.1 mM) (C) infected with HSV-1 (MOI=2).

(D) The mRNA of indicated inflammatory genes was determined by real-time PCR using the liver of wildtype, NAMPT-knockout and NAMPT-knockout mice with NR (400 mg/kg/day) at 3 days post-infection of HSV-1 (2x10$^7$ PFU, intravenous). ND, not detected. Related to Figure 2.

### Figure S4

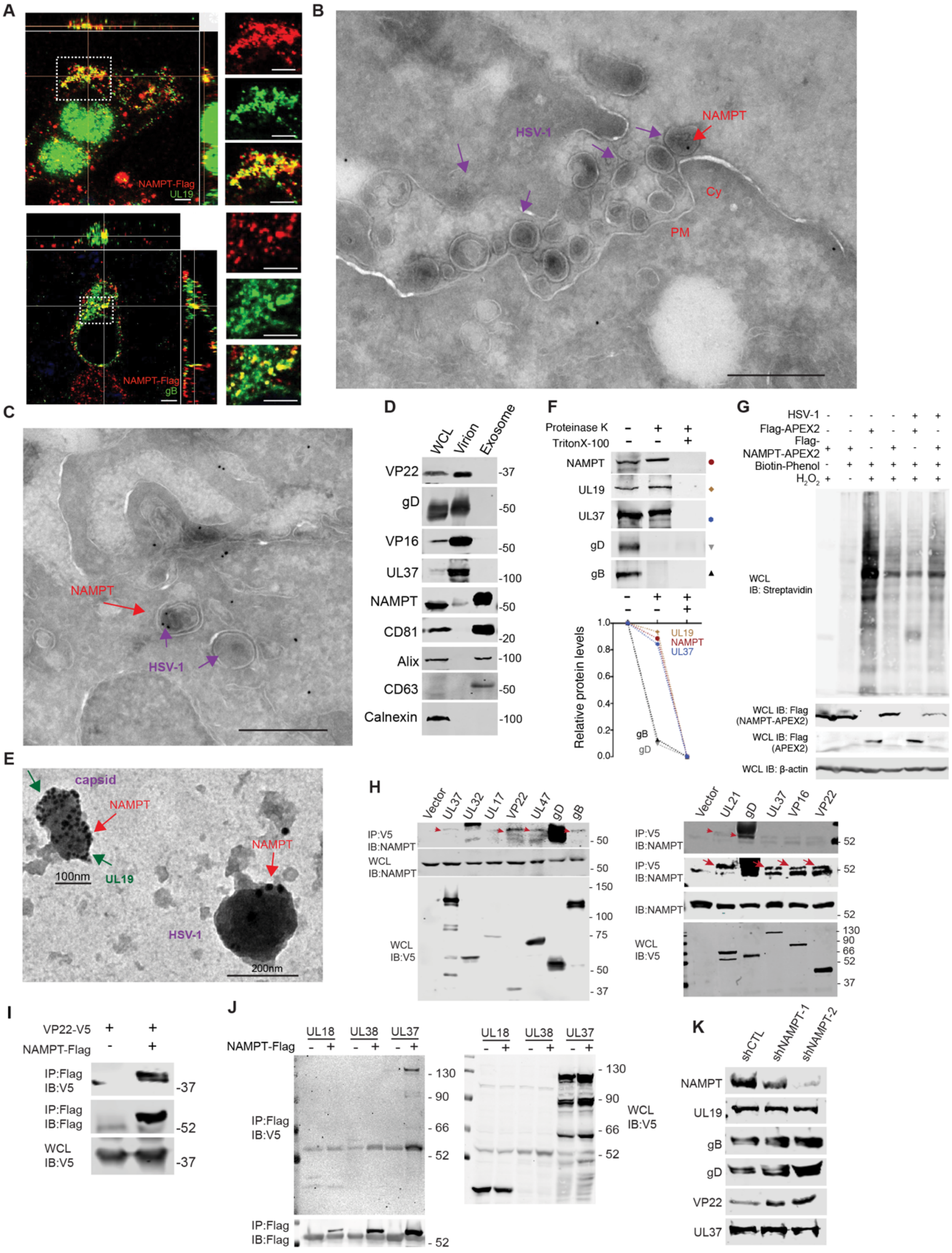

A
NAMPT-Flag
UL19
NAMPT-Flag
gB
B
HSV-1
NAMPT
Cy
PM
C
NAMPT
HSV-1
D
WCL
Virion
Exosome
VP22
gD
VP16
UL37
NAMPT
CD81
Alix
CD63
Calnexin
E
capsid
NAMPT
UL19
HSV-1
100nm
200nm
F
Proteinase K
TritonX-100
NAMPT
UL19
UL37
gD
gB
Relative protein levels
G
HSV-1
Flag-APEX2
Flag-NAMPT-APEX2
Biotin-Phenol
$H_2O_2$
WCL IB: Streptavidin
WCL IB: Flag (NAMPT-APEX2)
WCL IB: Flag (APEX2)
WCL IB: β-actin
H
Vector
UL37
UL32
UL17
VP22
UL47
gD
gB
IP:V5 IB:NAMPT
WCL IB:NAMPT
WCL IB:V5
UL21
VP16
IB:NAMPT
I
VP22-V5
NAMPT-Flag
IP:Flag IB:V5
IP:Flag IB:Flag
WCL IB:V5
J
UL18
UL38
UL37
NAMPT-Flag
IP:Flag IB:V5
IP:Flag IB:Flag
WCL IB:V5
K
shCTL
shNAMPT-1
shNAMPT-2
NAMPT
UL19
gB
gD
VP22
UL37

**Figure S4. NAMPT is incorporated into HSV-1 virions**

(A) Representative immunofluorescence images of HSV-1-infected sgNAMPT HeLa cells reconstituted with HA-NAMPT using antibodies to HA (NAMPT), UL19 and gB. Scale bars, 5 μm.

(B and C) Cryo-electron microscopy analysis of NAMPT in HSV-1-infected sgNAMPT HeLa cell reconstituted with HA-NAMPT. Scale bars, 500 nm.

(D) Purified HSV-1 virions, along with whole cell lysates and exosomes, were analyzed by immunoblotting with antibodies against known exosome markers and HSV-1 structural proteins.

(E) Transmission electron microscopy analysis of purified HSV-1 virions after permeabilization with 1% Triton X-100 and immunogold-staining with antibodies to UL19 (10 nm gold) and HA (NAMPT, 25 nm gold).

(F) Top panels: Immunoblotting analysis of purified HSV-1 virions treated with protease K, in the absence or presence of Triton X-100 (1%), with antibodies to NAMPT and viral proteins. Bottom panel: Diagram of sensitivity to proteinase K of HSV-1 envelope glycoproteins, tegument protein UL37 and capsid protein UL19 without or with Triton X-100 treatment. Proteins were indicated by symbols with distinct shapes and colors.

(G) Immunoblotting analysis of whole cell lysates (WCL) of HeLa cells stably expressing APEX or NAMPT-APEX and infected with HSV-1 for biotinylation assay, with $H_2O_2$ serves as a positive control.

(H) HSV-1 proteins, including UL21, gD, UL37, VP16, VP22, UL32, UL17, UL47, gB precipitated with endogenous NAMPT in transfected 293T cells.

(I and J) NAMPT interactions with VP22 (I), UL37, UL18 and UL38 (J) in transfected 293T cells were analyzed by co-immunoprecipitation and immunoblotting.

(K) Immunoblotting analysis of HSV-1 structural proteins with indicated antibodies using virions produced from shCTL and shNAMPT HepG2 cells that were normalized against the UL19 capsid protein.

Related to Figure 3.

### Figure S5

A

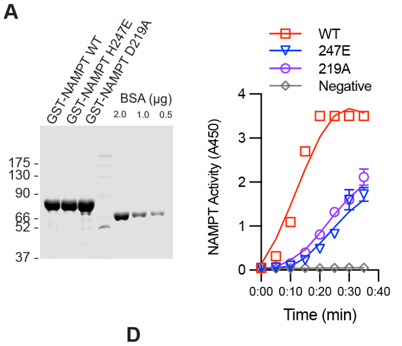


B

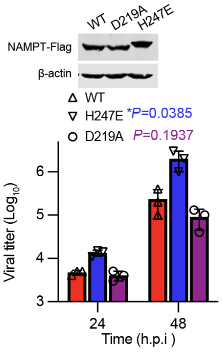


C

| | H247E | WT |
|---|---|---|
| Total pR peptide Abundance | $1.01\times10^{8}$ | $5.42\times10^{7}$ |
| Percentage (%) | 2.1 | 1.2 |

D

| Protein name | Peptide sequence | Positions |
|---|---|---|
| Tegument protein VP22 | NLLQR*ANE*LVNPDVVQDVDAATATR | R5 (254) and E8 (257) |
| | YGGSSSE*D*DEHPEVPR | E7 (74) and D8 (75) |
| | YGGSSSE*DDE*HPEVPR | E7 (74) and E10 (77) |
| | TDEDLNE*LLGITTIR (ADP-Ribosyl) | E7 (234) |
| | R*PVSGAVLSGPGPAR (ADP-Ribosyl) | R1 (86) |
| | SAQPE*SAALPDAPASTAPTR (ADP-Ribosyl) | E5 (149) |
| Envelope glycoprotein D | SETPNATQPELAPED*PEDSALL | D15 (297) |
| Envelope glycoprotein B | VASAAPSSPGTPGVAAATQAANGGPATPAPPAPGAPPTGD*PK | D40 (66) |
| Envelope glycoprotein H | D*TQPVAAIAPGFLAASAL | D1 (794) |
| Tegument protein UL21 | R*APR*PIAGSGAGSGGAGAKR | R1 (217), R4 (220) |
| Tegument protein VP16 | MDLLVD*E*LFAD*MNADGASPPPPRPAGGPK | D6 (6), E7 (7), D11 (11) |
| | SAATEE*PGAPLTTPPTLHGNQAR | E6 (314) |
| Tegument protein UL47 | LLMGD*EGAAAL | D5 (669) |
| | KDLE*PFPPAPVSGGSVLGPR | E4 (636) |
| Tegument protein UL46 | SAGPPPD*D*MASGPGGHR | D7 (439 ), D8 (440) |
| Inner tegument protein UL37 | AGPE*SPR | E4 (233) |
| Triplex capsid protein 2 UL18 | LADGFETD*IAIPSGISRPDAAALQR | D8 (9) |
| Capsid vertex component 1 UL17 | HPRPVLPPFDIAAEVAFTAR* | R20 (481) |
| | VAPAPHGPGGED*ILNDVLTLTPGTAKPR* | D12 (339), R28 (355) |
| | ATALTEACAAE*RVVR | E11 (433) |
| Major capsid protein UL19 | HRCLER*LIVETGSAVSTATAASDVQFK | R6 (1291) |
| Packaging protein UL32 | LAGTPAVWE*SGPQLR | E9 (353) |

E

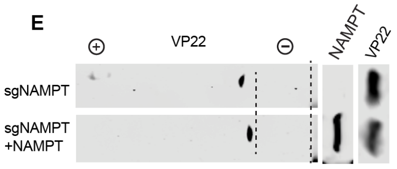


F

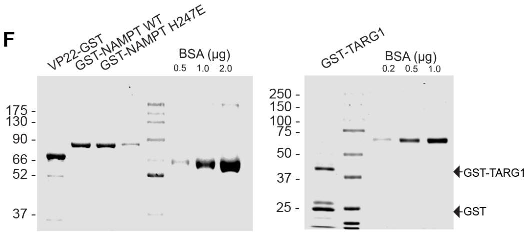


G

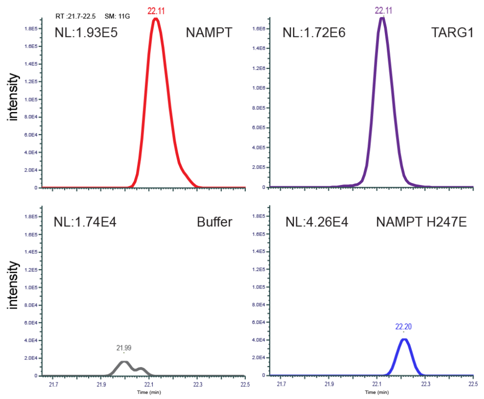


H

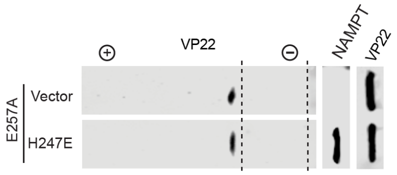


I

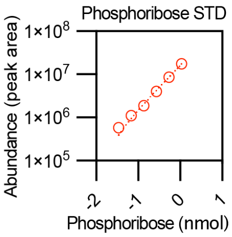

**Figure S5. NAMPT is a protein phosphoribosylase**

(A) Phosphoribosyltransferase activity in $NAD^+$ synthesis of NAMPT wildtype (WT), H247E and D219A mutants that were purified from bacteria. NAMPT proteins purified to high homogeneity were validated by Coomassie staining and shown on the left.

(B) HSV-1 titer in sgNAMPT HeLa cells reconstituted with NAMPT WT, H247E and D219A mutants at 24 and 48 h.p.i.

(C) Quantification of phosphoribosylated viral peptides in HeLa cells stably expressing wildtype NAMPT or the NAMPT-H247E mutant. Data represents one of three independent experiments.

(D) A list of identified phosphoribosylated and ADP-ribosylated (only in VP22) sites within their corresponding peptides, mapped to HSV-1 proteins. Numbers in parentheses indicate the corresponding residues within HSV-1 proteins.

(E) Two-dimensional gel electrophoresis and immunoblotting analysis of VP22 in sgNAMPT 293T cells and those reconstituted with NAMPT expression.

(F) Coomassie staining of purified GST-VP22, GST-NAMPT, GST-NAMPT-H247E (top panel) and GST-TARG1 (bottom panel).

(G) Detection of phosphoribose in reactions containing buffer, NAMPT, NAMPT-H247E, or TARG1 by LC-MS with GST-VP22 as the substrate.

(H) Two-dimensional gel electrophoresis and immunoblotting analysis of VP22-E257A in 293T cells transiently expressing the NAMPT-H247E mutant with antibody against the V5 epitope (VP22).

(I) Phosphoribose in serial dilutions was determined by LC-MS, which serves as a standard for phosphoribose released from the NAMPT phosphoribosylase reaction.

Statistical significance was calculated using unpaired two-tailed Student's *t*-tests.

Related to Figure 4.

### Figure S6

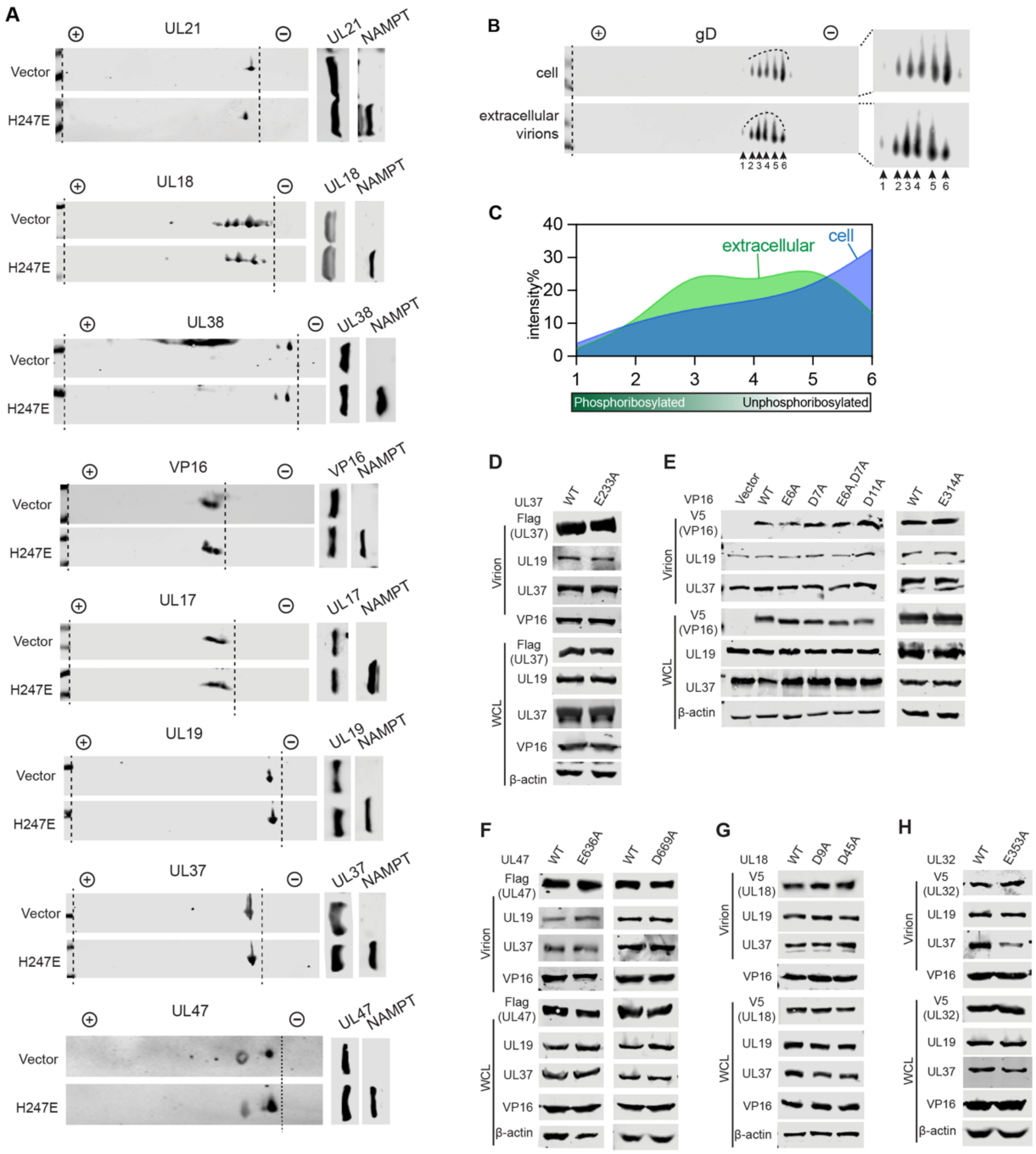
A
UL21
NAMPT
Vector
H247E
UL18
UL38
VP16
UL17
UL19
UL37
UL47
B
gD
cell
extracellular
virions
1 2 3 4 5 6
C
extracellular
cell
intensity%
40
30
20
10
0
1
2
3
4
5
6
Phosphoribosylated
Unphosphoribosylated
D
UL37
WT
E233A
Flag
(UL37)
UL19
UL37
VP16
β-actin
Virion
WCL
E
VP16
Vector
WT
E6A
D7A
E6A,D7A
D11A
E314A
V5
(VP16)
F
UL47
E636A
D669A
Flag
(UL47)
G
UL18
D9A
D45A
V5
(UL18)
H
UL32
E353A
V5
(UL32)

**Figure S6. Blockade of phosphoribosylation of structural proteins impedes their incorporation into HSV-1 virion**

(A) Whole cell lysates of 293T cells that were transiently transfected with plasmids containing the dominant negative NAMPT-H247E mutant and HSV-1 proteins, including UL21, UL18, UL38, VP16, UL17, UL19, UL37, UL47, were analyzed by two-dimensional gel electrophoresis and immunoblotting with antibodies to the V5 epitope (HSV-1 proteins) and FLAG (NAMPT).

(B and C) Two-dimensional gel electrophoresis and immunoblotting analysis of purified intracellular and extracellular HSV-1 virions (produced from sgNAMPT 293T cells) with antibodies to gD (B) and quantification by densitometry of gD species (C). Numbers at the bottom (B) and on x-axis (C) indicate the gD species with distinct charge status.

(D to H) Wild-type and phosphoribosylation-resistant mutants of HSV-1 proteins, including UL37 (D), VP16 (E), UL47 (F), UL18 (G) and UL32 (H), incorporated in extracellular HSV-1 virions and expressed in HSV-1-infected cells were analyzed by immunoblotting with indicated antibodies. All HSV-1 proteins were tagged with either the V5 epitope (VP16, UL18 AND ul32) or Flag epitope (UL37 and UL47).

Related to Figure 5.

Figure S7

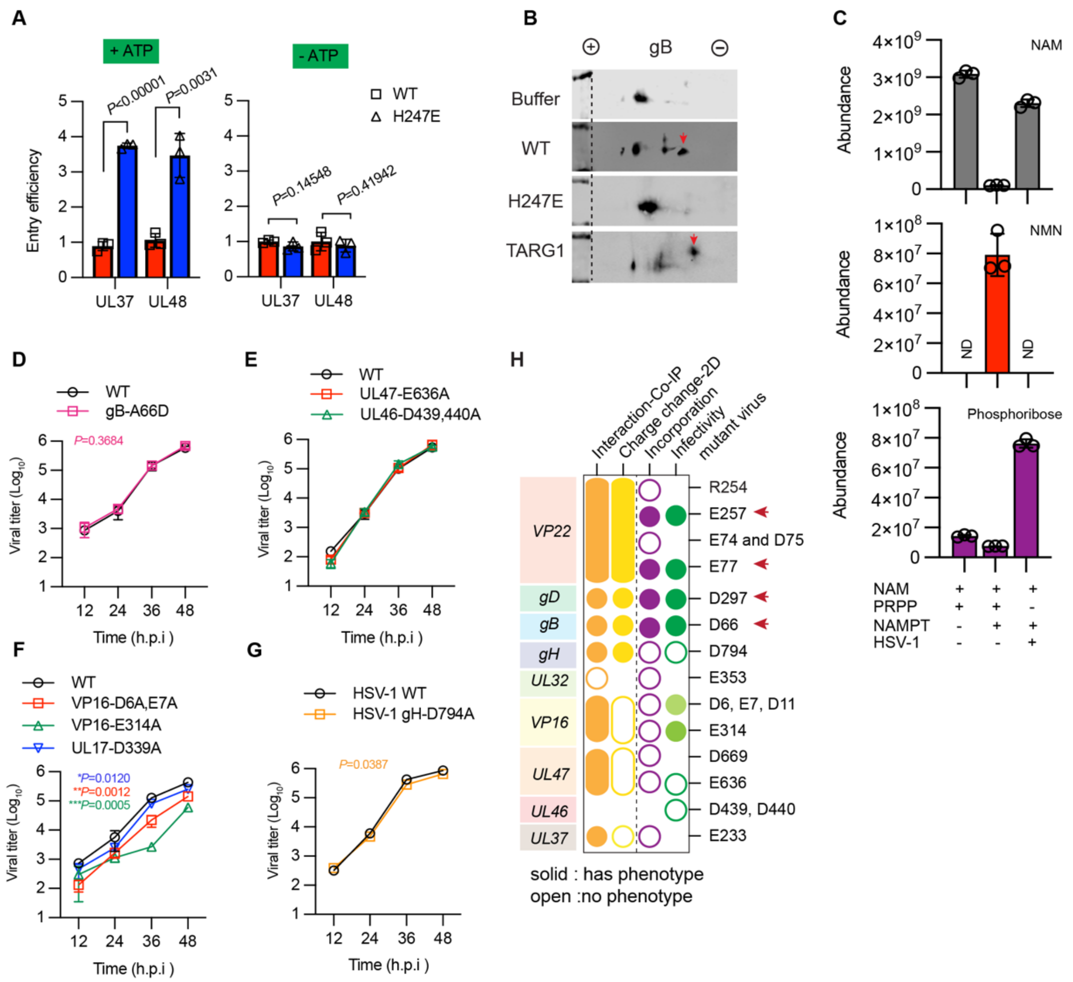


**Figure S7. Dephosphoribosylation of structural proteins impedes HSV-1 entry and replication**

(A) Entry analysis of HSV-1 virions treated with NAMPT or NAMPT-H247E, with or without ATP (2 mM).

(B) Two-dimensional gel electrophoresis and immunoblotting analysis of HSV-1 virions treated with wildtype NAMPT (WT), the NAMPT-H247E mutant or TARG1 with antibody against gB. Red arrows indicate the new species produced by NAMPT or TARG1 treatment.

(C) Quantification of NAM, NMN and phosphoribose in the supernatant of the in vitro phosphoribosylase reactions by mass spectrometry. ND, not detected.

(D) Growth curve of parental wildtype (WT) HSV-1 and recombinant HSV-1 carrying gB-A66D (revertant) in HepG2 cells (MOI=0.1) was determined by plaque assay using Vero cells.

(E to G) Growth curve of HSV-1 containing phosphoribosylation-resistant mutants of UL46 (D439, 440A), UL47 (E636A) (E), UL16 (D6A, E7A, D314A) and UL17 (D339A) (F) and gH (D794A) (G) in HepG2 cells (MOI=0.1) was determined by plaque assays using Vero cells.

(H) Summary of biochemical and functional characterization of the site-specific phosphoribosylation of HSV-1 proteins. Red arrows indicate the residues whose phosphoribosylation-resistant mutations displayed significant effect on HSV-1 infection. Intensity of color indicates the degree of effect and open circles indicate no phenotype. Statistical significance was calculated using unpaired two-tailed Student's *t*-tests and two-way ANOVA analysis.
Related to Figure 6.
